# CD4^+^ T cell lymphopenia and dysfunction in severe COVID-19 disease is autocrine TNF-α/TNFRI-dependent

**DOI:** 10.1101/2021.06.02.446831

**Authors:** Iulia Popescu, Mark E. Snyder, Carlo J. Iasella, Stefanie J. Hannan, Ritchie Koshy, Robin Burke, Antu Das, Mark J. Brown, Emily J. Lyons, Sophia C. Lieber, Xiaoping Chen, John C. Sembrat, Xiaojing An, Kelsey Linstrum, Georgios Kitsios, Ioannis Konstantinidis, Melissa Saul, Daniel J. Kass, Jonathan K. Alder, Bill B. Chen, Elizabeth A. Lendermon, Silpa Kilaru, Bruce Johnson, Matthew R. Morrell, Joseph M. Pilewski, Joseph E. Kiss, Alan H. Wells, Alison Morris, Bryan J. McVerry, Deborah K. McMahon, Darrell J. Triulzi, Kong Chen, Pablo G. Sanchez, John F. McDyer

**Affiliations:** Division of Pulmonary, Allergy, and Critical Care Medicine, Department of Medicine, University of Pittsburgh School of Medicine; Pittsburgh, Pennsylvania, 15213, USA; Department of Pharmacy and Therapeutics, University of Pittsburgh School of Pharmacy; Pittsburgh, Pennsylvania, 15213, USA; Department of Critical Care Medicine, University of Pittsburgh School of Medicine; Pittsburgh, Pennsylvania, 15213, USA; Department of Medicine, University of Pittsburgh School of Medicine, Pittsburgh; Pennsylvania, 15213, USA; Aging Institute, Department of Medicine, University of Pittsburgh School of Medicine; Pittsburgh, Pennsylvania, 15213, USA; Division of Transfusion Medicine, Department of Pathology, University of Pittsburgh School of Medicine; Pittsburgh, Pennsylvania, 15213, USA; Division of Laboratory Medicine, Department of Pathology, University of Pittsburgh School of Medicine; Pittsburgh, Pennsylvania, 15213, USA; Division of Infectious Diseases, Department of Medicine, University of Pittsburgh School of Medicine; Pittsburgh, Pennsylvania, 15213, USA; Department of Cardiothoracic Surgery, University of Pittsburgh School of Medicine; Pittsburgh, Pennsylvania, 15213, USA

## Abstract

Lymphopenia is common in severe COVID-19 disease, yet the mechanisms are poorly understood. In 148 patients with severe COVID-19, we found lymphopenia was associated with worse survival. CD4^+^ lymphopenia predominated, with lower CD4^+^/CD8^+^ ratios in severe COVID-19 compared to recovered, mild disease (p<0.0001). In severe disease, immunodominant CD4^+^ T cell responses to Spike-1(S1) produced increased *in vitro* TNF-α, but impaired proliferation and increased susceptibility to activation-induced cell death (AICD). CD4^+^TNF-α^+^ T cell responses inversely correlated with absolute CD4^+^ counts from severe COVID-19 patients (n=76; R=-0.744, P<0.0001). TNF-α blockade including infliximab or anti-TNFRI antibodies strikingly rescued S1-specific CD4^+^ proliferation and abrogated S1-AICD in severe COVID-19 patients (P<0.001). Single-cell RNAseq demonstrated downregulation of Type-1 cytokines and NFκB signaling in S1-stimulated CD4^+^ cells with infliximab treatment. Lung CD4^+^ T cells in severe COVID-19 were reduced and produced higher TNF-α versus PBMC. Together, our findings show COVID-19-associated CD4^+^ lymphopenia and dysfunction is autocrine TNF-α/TNFRI-dependent and therapies targeting TNF-α may be beneficial in severe COVID-19.

**One Sentence Summary:** Autocrine TNF-α/TNFRI regulates CD4^+^ T cell lymphopenia and dysfunction in severe COVID-19 disease.

## Main text

### INTRODUCTION

Severe viral pneumonia and respiratory disease due to SARS-CoV-2 infection has been the major clinical manifestation associated with patient mortality during the COVID-19 pandemic(*1–3*). The progression of upper respiratory symptoms to severe viral pneumonia, and at times the adult respiratory distress syndrome later in the course of infection in a subset of patients, have led many investigators to hypothesize an important role for inflammatory mediators or ‘cytokine storm’ in the course of disease, as multiple studies have shown increased systemic levels of cytokines during moderate and severe COVID-19 disease(*4, 5*). Treatment with the corticosteroids, dexamethasone and hydrocortisone, were found to reduce mortality in severe COVID-19 pneumonia and suggested that immune modulation could impact severe disease(*6, 7*). Several subsequent studies of IL-6 receptor antagonists in severe COVID pneumonia have found mixed results with some showing benefit(*8–10*), while others did not find efficacy(*11–13*).The recent NIH treatment guidelines recommend use of tocilizumab in severe COVID-19 with rapidly decompensating respiratory status in combination with corticosteroids https://www.covid19treatmentguidelines.nih.gov/statement-on-tocilizumab/. However, the role for immune modulators in severe COVID-19 disease remains incompletely defined, as ongoing studies have not yet been completed.

An early report correlated lymphopenia with poor outcomes in COVID-19 and disease severity(*14*). However, the mechanisms leading to lymphopenia in COVID-19 clinical syndrome remain poorly understood(*15*). Recent studies of the peripheral T cells during SARS-COV-2 infection have found an activated phenotype in both CD8^+^ and CD4^+^ T cells in severe disease, including increased surface expression of CD38^+^, CD95^+^, HLA-DR^+^, Ki67^+^ and programmed cell death protein 1 (PD-1), compared to mild disease and non-infected normal controls(*16, 17*). One study found that more profound lymphopenia was associated with increased serum levels of the inflammatory cytokines IL-6, IL-10 and TNF-α(*18*). Another study found that IL-6 levels negatively correlated with cytotoxic immune cells in severe COVID-19 disease(*19*). We hypothesized that factors such as T cell activation and the inflammatory milieu together, contributed to the development of COVID-19-associated lymphopenia during severe disease. We further reasoned that other T cell responses to SARS-CoV-2 proteins would be detected in mild and severe disease, in addition to spike, as an earlier studies have demonstrated a broad response to SARS-CoV-2 epitopes across CD4^+^ and CD8^+^ T cells(*20, 21*). We hypothesized that SARS-CoV-2-specific T cell responses might play an important role in the development of T cell lymphopenia, as in other viral infections such as HIV that result in activation-induced cell death (AICD)(*22*).

Here, we found that COVID-19-associated lymphopenia in severe disease is disproportionately a CD4^+^ T cell lymphopenia and is associated with increased mortality, with significantly reduced peripheral CD4^+^/CD8^+^ T cell ratios in severe disease compared to recently recovered mild COVID-19 patients. We further show that the immunodominant response of CD4^+^ T cells is S1-specific production of the pro-inflammatory Type-1 cytokine, TNF-α, in severe COVID-19 disease compared to control mild COVID-19 disease patients. We observed impaired CD4^+^ T cell proliferation and AICD via TNFRI signaling that could be rescued *in vitro* with various TNF-α blockade agents. Similarly, reduced CD4^+^ numbers and S1-specific TNF-α predominant responses were detected at higher frequencies in resident lung T cells from patients with recent severe COVID-19 pneumonia. Together, our findings show that CD4^+^ T cell lymphopenia and dysfunction in severe COVID-19 respiratory disease is autocrine CD4^+^ TNF-α/TNFRI-dependent.

### RESULTS

#### Peripheral lymphopenia is associated with mortality, with a predominant CD4^+^ T cell lymphopenia in severe COVID-19 disease

We hypothesized that peripheral lymphopenia was associated with poor outcomes in patients with severe COVID-19 disease. We evaluated a multi-hospital cohort (n=148) within our medical system of severe patients hospitalized with documented SARS-CoV-2 infection, with demographic data shown in Table S1. We evaluated the first absolute lymphocyte count (ALC) on hospitalization and found the median to be 700/mm^3^ (Fig. 1A). We next assessed 30-day mortality by Kaplan-Meier and observed increased mortality in those below the median (Fig. 1B). We performed flow cytometry on isolated PBMC from both groups (n=76 total; gating strategy Fig. S1A) and observed a significant diminution in CD4^+^ T cell frequencies and reduced CD4^+^/CD8^+^ T cell ratios in patients with lymphopenia (Fig. 1C-E). We observed significantly reduced absolute CD4^+^ and CD8^+^ counts (though less profound) (Fig. 1F, G), based on ALC (Fig. S1B). Together, these data indicate that COVID-19-associated lymphopenia is a predominantly CD4^+^ T cell lymphopenia. To further evaluate T cell function and phenotype, we randomly selected a sub-cohort of 24 severe COVID-19 patients, of which n=9 had lymphopenia (Fig. S1C), and a control cohort (n=24) of mild COVID disease (Table 1) (Fig. 1H), immediately after resolution of symptoms and undergoing evaluation for potential plasma donation. While blood counts were not available for mild, recovered COVID patients we observed significantly higher CD4^+^/CD8^+^ ratios compared to our severe COVID-19 sub-cohort (Fig. 1H).

**Figure 1.**
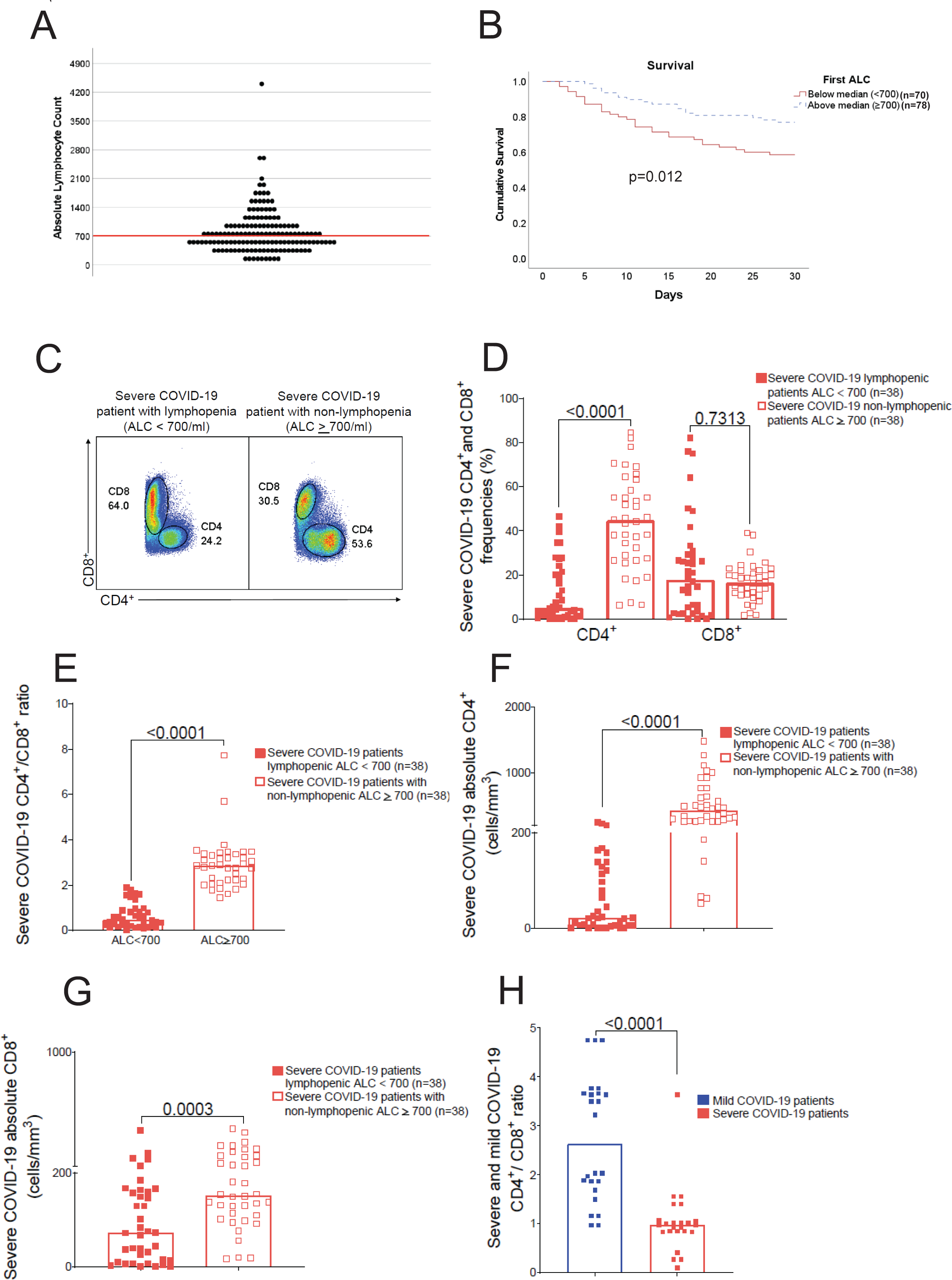
Peripheral lymphopenia is associated with mortality and a predominant CD4^+^ T cell lymphopenia in severe COVID-19 disease. *(A)* The distribution of absolute lymphocyte counts (ALC) for n=148 severe COVID-19 patients. Values represent the first ALC obtained upon admission and the median ALC *(red line)* was 700 cells/mm^3^; *(B)* Kaplan-Meier 30-day survival curves severe COVID-19 patients with initial ALC values at or above the median (n=78) vs. those below the median (n=70), showing a significant difference in survival between groups (Log-rank test p=0.012). *(C)* Representative flow cytometry plot of CD4^+^and CD8^+^ T cells from severe COVID-19 patients with lymphopenia ALC<700 *(left panel)* and non-lymphopenia ALC≥700 *(right panel)* with frequencies of T cell subsets. *(D)* Cumulative data (n=76) from severe COVID-19 patients showing frequencies of CD4^+^ and CD8^+^ T cell subsets with lymphopenia (ALC<700) (n=38) *(full red squares)* compared with non-lymphopenia (ALC≥700) patients (n=38) *(empty red squares)*. *(E)(F)(G)* Pooled data (n=76) from severe COVID-19 patients showing CD4^+^/CD8^+^ T cells ratio *(E)* the absolute CD4^+^ T cells numbers (cells/mm^3^) *(F)* and the absolute CD8^+^ T cells numbers (cells/mm^3^) *(G)* with lymphopenia (ALC<700) (n=38) *(full red squares)* compared with non-lymphopenia (ALC≥ 700) patients (n=38) *(empty red squares)*. *(H)* Cumulative data (n=48) from the cohort used in functional studies COVID-19 patients CD4^+^/CD8^+^ T cells ratio with mild disease (n=24) *(blue square)* compared with severe disease patients (n=24) *(red square)*. Bars represent median values, and P values were calculated using the Mann-Whitney-Wilcoxon test. Statistical analysis performed using Mann-Whitney U-test with a two-sided p value of < 0.05 considered statistically significant.

**TABLE 1.**

|  | **Severe (n = 24)** | **Mild (n = 24)** | **p-value** |
| --- | --- | --- | --- |
| **Age, median [IQR]** | 69 [55-73] | 45 [33-64.5] | <0.01 |
| **Race** |  |  | 0.11 |
| White | 15 (62.5%) | 19 (79.2%) |  |
| Black | 5 (20.8%) | 2 (8.3%) |  |
| Other | - | 3 (12.5%) |  |
| Unknown/No Answer | 4 (16.7%) | - |  |
| **Female** | 11 (45.8%) | 10 (41.7%) | 0.77 |
| **Peak Resp Support** |  |  |  |
| Low flow NC | 1 (4.2%) | - |  |
| High low NC | 5 (20.8%) | - |  |
| NIPPV | 1 (4.2%) | - |  |
| Mechanical Ventilation | 16 (66.7%) | - |  |
| ECMO | 1 (4.2%) | - |  |
| **Cardiovascular Disease** | 11 (45.8%) | 7 (29.2%) | 0.23 |
| **Respiratory Disease** | 2 (8.3%) | 5 (20.8%) | 0.42 |
| **Diabetes** | 9 (37.5%) | 1 (4.2%) | 0.01 |

NC – nasal cannula; NIPPV – non-invasive partial pressure ventilation; ECMO – extracorporeal membrane oxygenation

#### CD4^+^TNF-α^+^ and CD4*^+^*CD107a^+^ Spike-1-specific effector responses are immunodominant and increased in severe COVID-19-infected patients

We next assessed COVID-specific and superantigen-induced T cell effector responses (*Staph.* Enterotoxin B-SEB) in our mixed COVID-19 cohort (mild and severe disease; total n=48). Using pooled peptides to the major SARS-CoV-2 antigens we assessed both antigenic and effector immunodominance (major T cell response) in PBMC following a 6 h *in vitro* re-stimulation. Spike-1 (S1) responses (TNF-α and IFN-γ) were found to be immunodominant in CD4^+^ T cells compared to the other antigens (Spike-2 (S2), viral envelope membrane protein (VEMP) and nucleocapsid (NCAP)) (Fig. 2A-C). A similar immunodominance pattern was found for COVID-specific CD8^+^IFN-γ^+^ T cell responses (Fig. S2A). We further compared blood S1-specific effector responses between CD4^+^ and CD8^+^ T cells in our severe COVID-19 cohort and found that S1-specific TNF-α responses predominated in CD4^+^ T cells, in contrast to S1-specific IFN-γ^+^ responses in CD8^+^ T cells, but with similar frequencies of cytotoxic CD107a^+^ responses (Fig. 2D). The overall hierarchy of T cell effector responses were found to be predominant Type-1 immune responses, with little IL-17a or IL-13 COVID S1-specific responses detected. We next determined whether there were differences adjusting for COVID-19 disease severity and found that S1-specific CD4^+^ responses remained overall immunodominant and with higher frequencies in severe disease (Fig. S2B), with CD4^+^TNF-α^+^ and CD4^+^CD107a^+^ responses in particular increased among severe COVID-19 patients compared to mild disease (Fig. 2E). One exception was S1-specific CD4^+^IL-2^+^ frequencies which were significantly reduced among severe patients compared with mild disease patients. Notably, the hierarchy of CD4^+^ TNF-α^+^ > IFN-γ^+^ responses in severe disease was observed with S1, and VEMP to a lesser extent, but not to other COVID antigens (Fig. S2C). Conversely, COVID-specific CD8^+^IFN-γ^+^ > CD8^+^TNF-α^+^ cell responses were found to be significantly increased among those with severe versus mild disease, whereas CD8^+^CD107a^+^ responses did not differ (Fig. 2F). We next compared the multifunctionality of S1-specific CD4^+^ versus CD8^+^ effector T cell responses. We found that CD4^+^ T cell multifunctional responses were significantly reduced in severe COVID disease compared to CD8^+^ cells, and with more single^+^ CD4^+^TNF-α^+^ frequencies (Fig. 2G). CD4^+^ T cells demonstrated a significant reduced multifunctionality in severe versus mild COVID-19 patients, with reduced IL-2 and increased CD4^+^TNF-α^+^ and CD4^+^CD107a^+^ frequencies (Fig. 2H), in contrast to CD8^+^ T cell multi-functionality, which did not differ based on COVID-19 disease severity (Fig. S2D). We found that S1-specific CD4^+^TNF-α^+^ responses frequencies were inversely correlated with CD4^+^/CD8^+^ ratios in our mixed COVID-19 cohort (n=48) (Fig. 2I). Next, we measured S1-specific CD4^+^TNF-α^+^ responses in our expanded number of severe COVID patients (n=76) and found this to be an inverse immune correlate of CD4^+^ lymphopenia (Fig. 2J) and CD4^+^/CD8^+^ ratios (Fig. S2E). In contrast, S1-specific CD4^+^IFN-γ^+^ or CD4^+^CD107a^+^ responses were not as strong an immune correlate with CD4^+^ numbers. Taken together, COVID-specific CD4^+^ T cells are skewed in severe COVID-19 disease to produce high levels of TNF-α that inversely correlate with absolute CD4^+^ counts.

**Figure 2.**
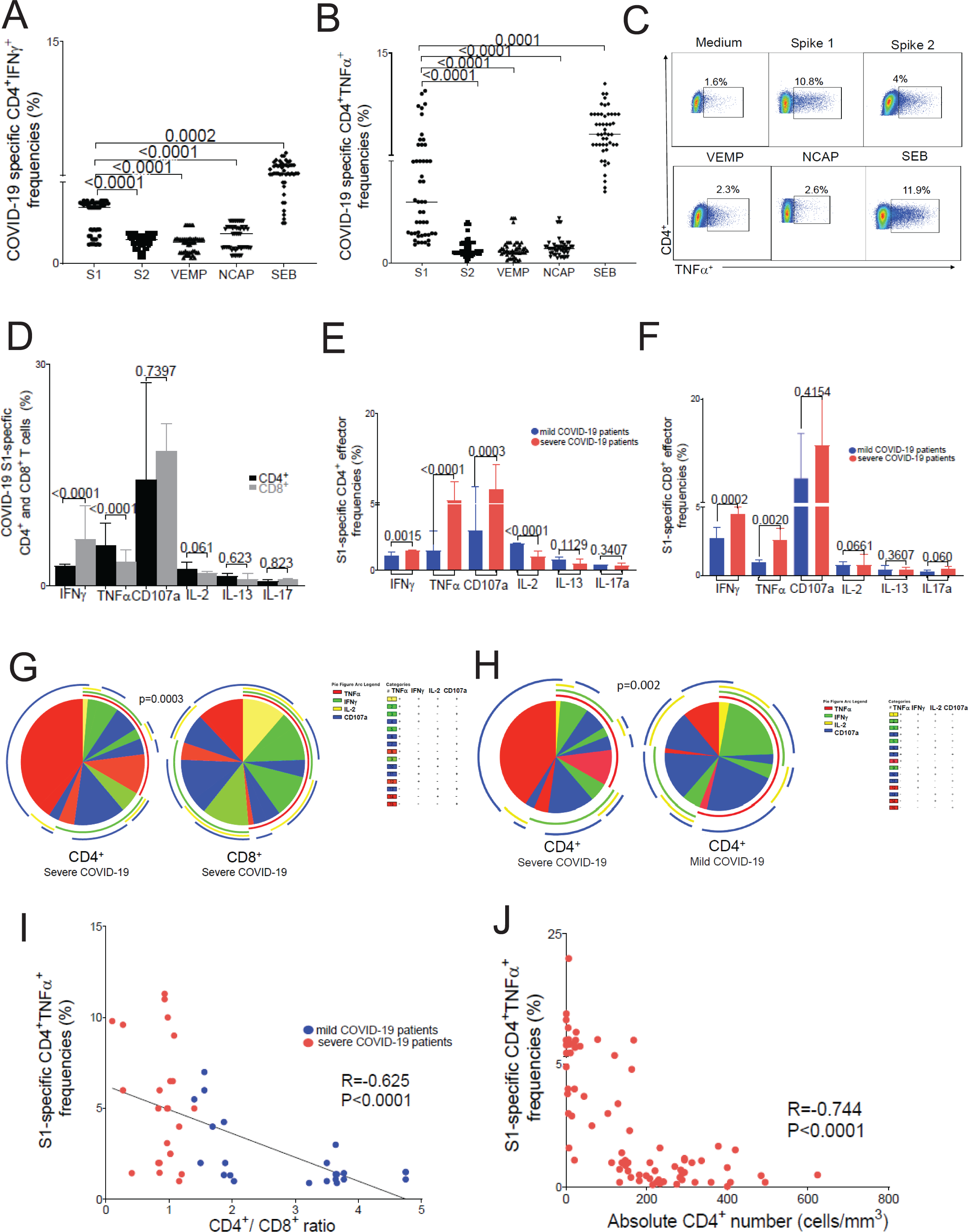
CD4^+^TNF-α^+^ and CD4*^+^*CD107a^+^ Spike-1-specific effector responses are immunodominant and increased in severe COVID-19-infected patients. Pooled data of intracellular cytokine staining immunodominance in total COVID-19 cohort (n=48) for Spike 1 (S1), Spike 2 (S2), viral envelope small membrane protein (VEMP) and nucleocapsid (NCAP) COVID-19 specific and SEB-reactive CD4^+^ T-cell effector responses. Values represent frequencies following 6h *in vitro* stimulation with respective pooled peptides or SEB minus medium alone (unstimulated) of CD4^+^IFNγ^+^ *(A)* and CD4^+^TNFα^+^ *(B). (C)* Representative flow cytometry plots data of intracellular cytokine staining of severe COVID-19 CD4^+^TNFα^+^ specific S1, S2, *(upper panels)* VEMP, NCAP and SEB *(lower panels)* responses and medium alone *(upper left panel)*. *(D)* Cumulative data of intracellular staining of Spike 1 (S1)-specific COVID-19 CD4^+^ *(black columns)* and CD8^+^ *(grey columns)* T-cell effector responses (IFNγ, TNFα, CD107a, IL-2, IL-13, IL-17) in severe COVID-19 cohort. Bars represent median values, and *P* values were calculated using the Mann-Whitney-Wilcoxon test. Statistical analysis performed using Mann-Whitney U-test with a two-sided p value of < 0.05 considered statistically significant. Pooled data showing intracellular staining of S1-specific COVID-19 CD4^+^ *(E)* and CD8^+^ *(F)* T-cell effector responses (IFNγ, TNFα, CD107a, IL-2, IL-13, IL-17) in mild *(blue columns)* vs. those with severe *(red columns)* COVID-19. Bars represent median values, and *P* values were calculated using the Mann-Whitney-Wilcoxon test. Statistical analysis performed using Mann-Whitney U-test with a two-sided p value of < 0.05 considered statistically significant. Individual pie charts showing CD4^+^ *(left pie)* and CD8^+^ *(right pie)* S1-specific (n=24) multifunctional responses in severe COVID-19 *(G)* and pie charts showing CD4^+^ in severe *(left pie)* vs. CD4^+^ in mild COVID-19 disease *(right pie)* T-cell multifunctional responses (n=48) *(H)*. The T-cell subset multifunctional responses are for four cytokines: TNFα (*red arch*), IFNγ (*green arch*), IL-2 (*yellow arch*) and CD107a (*blue arch*) and the combination of cytokine responses: +4-*yellow pie fraction*; +3-*green pie fraction;* +2-*blue pie fraction* and +1-*red pie fraction*, using Boolean analysis. Significant differences when comparing mean frequencies of single and multifunctional responses are indicated by a p *≤* 0.05. All p-values were determined by the Kruskal-Wallis one-way ANOVA or Wilcoxon signed-rank test. Data analyzed using the program SPICE. *(I)* Inverse correlation of S1-specific COVID-19 CD4^+^TNFα^+^ response frequencies with CD4^+^/CD8^+^ ratio on severe *(red dots)* and mild *(blue dots)* COVID-19 patients n=48. *(J)* The inverse correlation of S1-specific COVID-19 CD4^+^TNFα^+^ response frequencies with absolute CD4^+^ number (cells/mm^3^) on n=76 severe COVID-19 *(red dots).* Analysis was performed using the Spearman rho test and correlation coefficient R and P values were calculated using Spearman rank correlation test.

#### Impaired S1-specific CD4^+^ T cell proliferation in severe COVID-19 correlates with lymphopenia, is autocrine TNF-α/TNFRI-dependent and can be rescued *in vitro* by infliximab

Because we observed CD4^+^ T cell lymphopenia and reduced CD4^+^/CD8^+^ ratios among severe COVID-19 patients, we hypothesized that COVID-specific CD4^+^ T cell proliferative capacities contributed to low CD4^+^ numbers. Therefore, we assessed S1-specific CD4^+^ T cell *in vitro* proliferation at 6 days following peptide re-stimulation and CFSE dilution and found significantly impaired proliferative responses in severe COVID patients compared with mild disease (Fig. 3A, B). While S1-specific CD8^+^ T cells proliferative responses were also comparatively reduced in the severe disease cohort versus the mild disease cohort, impaired CD4^+^ proliferation was more pronounced in severe disease, including in response to SEB (Fig. S3A-D). As we detected reduced S1-specific IL-2 production from CD4^+^ T cells in severe COVID-19, we determined whether exogenous IL-2 could rescue impaired proliferative responses. The addition of IL-2 to cultures significantly rescued S1-specific CD4^+^ T cell proliferation (Fig. S3E, F) as well as enhanced S1-specific CD8^+^ proliferative responses (Fig S3G), suggesting a relative IL-2-deficient state in severe COVID-19. We next determined that CD4^+^ proliferative responses correlated with CD4^+^/CD8^+^ ratios in the severe/mild COVID-19 cohort and within the severe disease cohort correlated with the absolute lymphocyte count (Fig. 3C, D). Taken together, our data support impaired CD4^+^ T cell proliferation as a mechanism for decreased CD4^+^ counts in severe COVID-19 disease.

**Figure 3.**
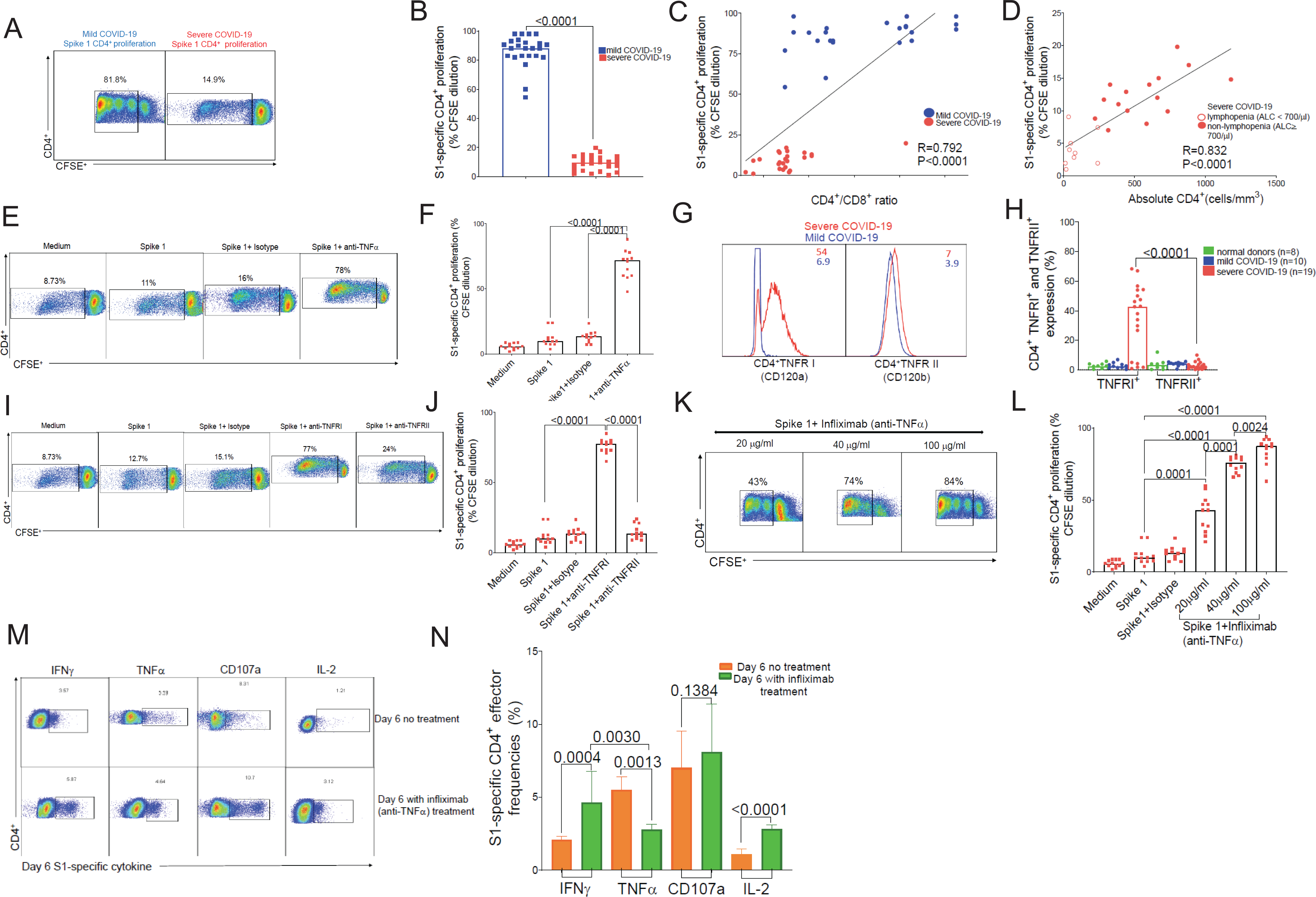
Impaired S1-specific CD4^+^ T cell proliferation in severe COVID-19 correlates with lymphopenia, is autocrine TNF-α/TNFRI-dependent, and can be rescued *in vitro* by infliximab. Representative flow cytometric plots *(A)* and cumulative data *(B)* showing day-6 S1-specific CD4^+^ T cell proliferation by CFSE dilution S1-specific from mild COVID-19 *(left panel-A and blue square-B)* vs. severe *(right panel-A and red square-B)* patients. Numbers indicate frequencies of live-gated CFSE diluted events (%) and bars represent median values and p-values (Mann-Whitney-Wilcoxon test). *(C)* Direct correlation of day-6 S1-specific COVID-19 CD4^+^ T-cell proliferation response frequencies with CD4^+^/CD8^+^ ratio from mild *(blue dots)* and severe COVID-19 *(red dots)*, n=48 patients. Analysis was performed using the Spearman rho test and correlation coefficient (R) and (P) values were calculated using Spearman rank correlation test. *(D)* Correlation of day-6 S1-specific COVID-19 CD4^+^ T-cell proliferation response frequencies with absolute CD4^+^ numbers in severe COVID-19 with lymphopenia *(empty red dots)* and non-lymphopenia *(full red dots),* n=24 patients, using Spearman rho and Spearman rank correlation tests. Representative flow cytometric plots *(E)* and cumulative data *(F)* showing day-6 S1-specific CD4^+^ T cell proliferation from CD8^+^-depleted PBMC by CFSE dilution S1-specific COVID-19 in severe disease patients in the presence or absence of anti-TNFα antibodies (20 μg/ml), n=12*. (G)* Representative histograms plots showing CD4^+^ T cell TNFRI^+^ (CD120a) *(left panel)* and TNFRII^+^ (CD120b) *(right panel)* surface expression in mild COVID-19 *(blue lines)* overlayed with severe COVID-19 *(red lines) (G).* Cumulative data *(H)* showing CD4^+^ T cell TNFRI^+^ (CD120a) and TNFRII^+^ (CD120b) surface expression in mild COVID-19 (n=10) *(blue dots)* vs. severe COVID-19 (n=19) *(red dots)* vs. normal donors (n=8) *(green dots)*. Representative flow cytometric plots *(I)* and cumulative data *(J)* of day-6 CD4^+^ S1-specific proliferation by CFSE dilution in the presence or absence of anti-TNFRI or anti-TNFRII Abs (20 μg/ml) in CD8^+^- depleted PBMC from severe COVID-19 *(red squares)* patients (n=12). Representative flow cytometric plots *(K)* and cumulative data *(L)* showing day-6 CD4^+^ S1-specific proliferation in the presence or absence of infliximab at various doses *(red squares)*. Bars represent median values and *p*-values were calculated using the Mann-Whitney-Wilcoxon test. Representative flow cytometric plots *(M)* and cumulative data *(N)* showing day-6 S1-specific frequencies of CD4^+^T cell effector responses (TNFα, IFNγ, CD107a and IL-2) following 6h re-stimulation with S1 peptides with *(lower panel -M, green columns-N)* or without *(upper panels -M, orange columns - N)* infliximab (100 μg/ml). Bars represent median values ± interquartile range for S1-specific CD4^+^ frequencies at day-6 *(orange columns)* without vs. day-6 with infliximab treatment *(green columns)* and *p*-values were calculated using the paired-Wilcoxon test with a two-sided p value of < 0.05 considered statistically significant.

We next hypothesized that increased S1-specific TNF-α production from *autologous* CD4^+^ T cells negatively impacted proliferative capacities and subsequently, CD4^+^ T cell numbers. To test this, we assessed the effect of TNF-α blockade on S1-specific CD4^+^ proliferation at 6 days in PBMC that were CD8^+^ T cell-depleted (96.2% purity; Fig. S3H) and found that TNF-α blockade resulted in significant restoration of CD4^+^ proliferative responses (Fig. 3E, F). We concomitantly measured TNF-α production from monocytes in the same cultures and found low TNF-α^+^ frequencies following peptide re-stimulation (Fig. S3I, J). Together, these data support autocrine CD4^+^ T cell-derived TNF-α as a significant source contributing to impaired S1-specific proliferation in severe COVID-19 disease. Similarly, in the initial set of proliferation experiments in which CD8^+^ T cells were not depleted, we observed enhancement of S1-specific CD8^+^ proliferation in the presence of TNF-α blockade (Fig. S3K). Next, we assessed TNFRI (CD120a) and TNFRII (CD120b) receptor surface expression in CD4^+^ T cells across our cohort and found TNFRI surface expression was strikingly up-regulated in CD4^+^ T cells from patients with severe disease compared to mild disease controls and normal donors (Fig. 3G, H). Targeted blockade of TNFRI resulted in marked restoration of S1-specific CD4^+^ proliferative responses, similar to TNF-α blockade, whereas TNFRII blockade had no significant effect (Fig. 3I, J). Finally, we tested the impact of the anti-TNF-α therapeutic, infliximab, and found it capable of significantly rescuing S1-specific CD4^+^ proliferation in a dose-dependent manner (Fig. 3K, L). Moreover, infliximab *in vitro* treatment resulted in enhanced CD4^+^ IFN-γ, IL-2, CD107a and reduced TNF-α responses at 6 days following secondary S1-peptide re-stimulation, thus reversing the hierarchical dominance of TNF-α in the Type-1 response (Fig. 3M, N). Together, our findings show autocrine CD4^+^ TNF-α/TNFRI-dependent regulation of impaired CD4^+^ proliferation and altered effector cytokines in severe COVID disease that can be reversed *in vitro* by infliximab.

#### Spike-1 induces activation-induced cell death in CD4^+^ T cells from severe disease patients and is autocrine TNF-α-/TNFRI-dependent

We further hypothesized that high COVID-specific TNF-α secretion from CD4^+^ T cells might contribute to activation induced cell death (AICD) and apoptosis, contributing to CD4^+^ T cell lymphopenia. To test this, we evaluated CD4^+^ T cells for Annexin V expression in short-term cultures (18 h) following re-stimulation *in vitro* with S1 peptides, in CD8^+^-depleted cultures. We found that Annexin V induction was significantly increased in severe COVID compared to mild disease patients (Fig. 4A, B). The addition of anti-TNF-α antibodies, including anti-TNFRI antibodies significantly inhibited Annexin V induction, as well as infliximab in a dose-dependent manner, but not anti-TNFRII antibodies (Fig. 4C, D). We also found that anti-Fas neutralizing antibodies inhibited Annexin V induction in S1-activated CD4^+^ T cells, but to a lesser extent than TNF-α blockade (Fig. S4A, B). Induction of AICD in cells from patients with severe disease occurred in the setting of other markers of activation, such as CD95, CD38 and to a lesser extent PD-1 compared to mild disease or normal controls (Fig. S4A). We observed that surface TNFRI expression correlated with surface CD38 expression (Fig. S4B). Thus, S1-induced CD4^+^ T cell AICD in severe COVID disease is a predominantly autocrine TNF-α/TNFRI-dependent mechanism contributing to CD4^+^ lymphopenia.

**Figure 4.**
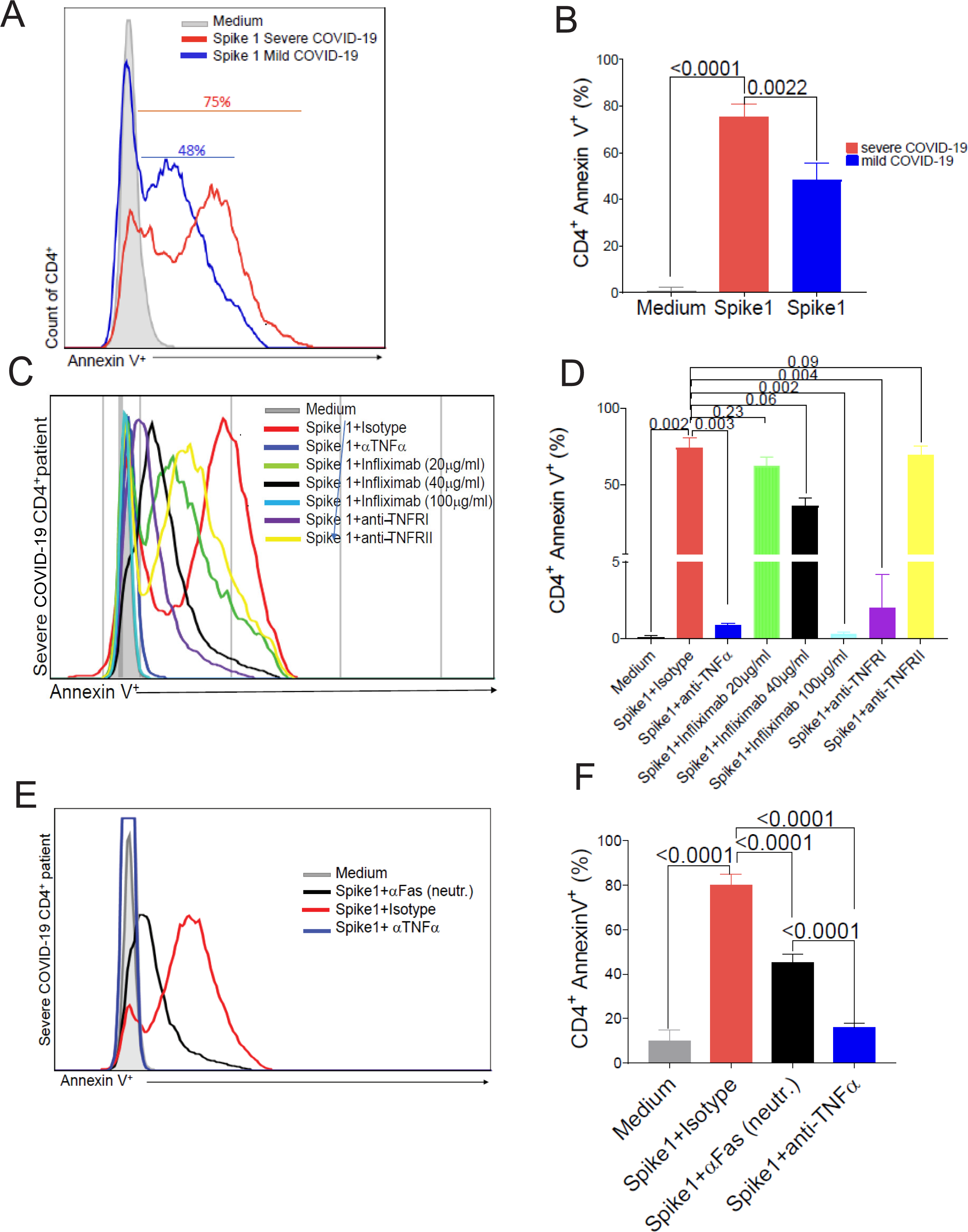
Spike-1 induces activation-induced cell death in CD4^+^ T cells from severe disease patients and is autocrine TNF-α-/TNFRI-dependent. *(A)* Representative histograms overlayed depicting the Annexin V^+^ staining in CD4^+^ T cells from severe *(red line)* vs mild *(blue line)* COVID-19 and medium as control *(grey full line)*. *(B)* Pooled data showing CD4^+^Annexin V^+^ in severe *(red column)* vs. mild *(blue column)* COVID-19 disease. Representative histograms *(C)* and pooled data *(D)* depicting the annexin V^+^ CD4^+^ in CD8^+^-depleted PBMC from severe COVID-19 patients. Cells were cultured in the presence or absence of anti-TNFα neutralizing antibodies (20 μg/ml) *(dark blue histogram lines-C and dark blue column-D)*, infliximab at various doses *(green,black and light blue histogram lines-C and columns-D)*, anti-TNFRI *(purple histogram lines-C and purple column-D)* or anti-TNFRII *(yellow histogram lines-C and yellow column-D)* antibodies (20 μg/ml), compared to S1-specific+isotype control (IgG1) *(red histogram lines-C and red column-D)*) or medium alone *(grey full histogram lines-C and grey column-D).* Bars represent cumulative median frequencies CD4^+^ Annexin V^+^ cells and P values were calculated using the Mann-Whitney test. Representative histograms *(E)* and pooled data *(F)* depicting the Annexin V^+^ CD4^+^ from COVID-19 severe patients in the presence or absence of anti-Fas neutralizing antibodies, compared to anti-TNFα neutralizing antibodies vs. S1-specific + isotype control (IgG1) or medium alone. Bars represent cumulative median frequencies ± interquartile range CD4^+^ AnnexinV^+^ cells from severe COVID-19 patients. P values were calculated using the Mann-Whitney test.

#### Single cell RNA sequencing of Spike-1-stimulated CD4^+^ T cells reveals striking TNF-α-dependent activation blocked by infliximab

We next examined the impact of TNF-α blockade on CD4^+^ T cell gene expression by performing single cell RNA sequencing of peripheral CD4^+^ T cells from two patients with severe COVID stimulated *in vitro* with S1-peptides in the presence or absence of infliximab. Following negative selection bead isolation of CD4^+^ T cells, the single cell RNAseq transcriptome identified 7 distinct clusters (Fig 5A), largely differentiated by the presence or absence of anti-TNF-α blockade (Fig 5B, C). Those CD4^+^ T cells exposed to S1-stimulation in the presence of infliximab demonstrated down-regulation of genes related to pro-inflammatory cytokines (*IL2*, *IFNG*, *TNFa*), co-stimulation (*TNFSF14* (*LIGHT)* and *TNFSF5* (*CD40LG*), NF-Kappaβ signaling pathway (*NFKBID*), antigen binding and activation (*SLAMF1*), and apoptosis (*FASLG, MYC, BCL2A1, SELENOK, NR4A1*) (Fig 5D, E). Gene set enrichment analysis of Hallmark gene sets identified upregulation of TNF SIGNALING VIA NFKB, ALLOGRAFT REJECTION, and IL2-STAT5 SIGNALING pathways among the CD4^+^ T cells stimulated with S1 alone (Fig. 5D). Together, these data show that TNF-α blockade using infliximab has a profound effect on S1-stimulated CD4^+^ T cells from severe COVID-19 patients.

**Figure 5.**
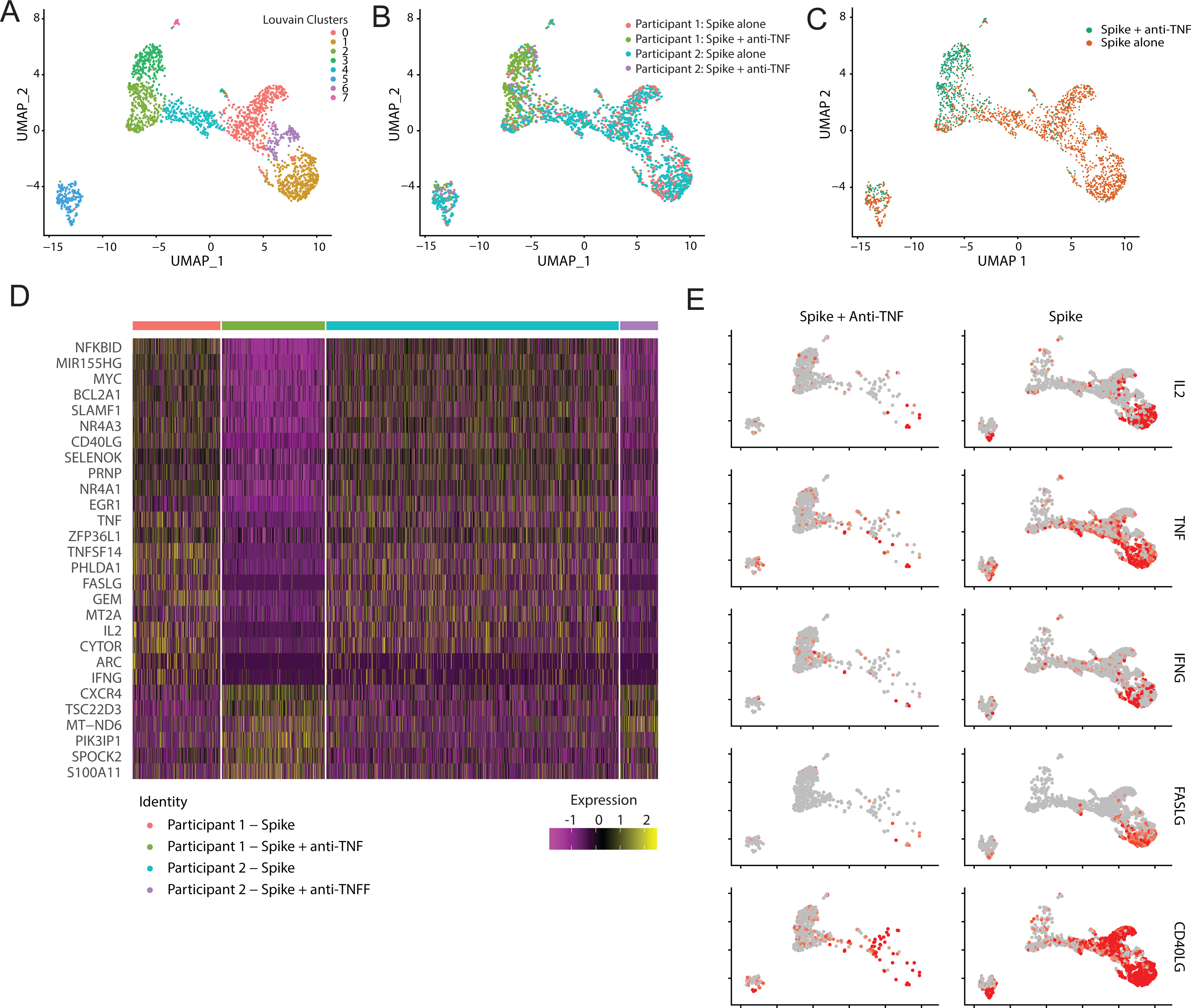
Single cell RNA sequencing of Spike-1-stimulated CD4^+^ T cells reveals striking TNF-α-dependent activation blocked by infliximab. Single cell RNA sequencing from bead isolated peripheral CD4^+^ T cells by negative selection 3 h following S1-stimulation of whole PBMC in the presence or absence of infliximab. *(A)* Uniform manifold approximation and projection (UMAP) of unsupervised clusters of cells with elevated CD3E gene expression and negative CD68, CD19, and CD8A gene expression; *(B)* Clusters by hash tag oligos (HTO); *(C)* clusters by experimental condition, spike protein alone versus spike protein and anti-TNF therapy in vitro; *(D)* Heatmap of highly variable genes by HTO; *(E)* Feature plot for select transcripts with red indicating upregulated cells.

#### High TNF-α production by lung resident memory CD4^+^ T cells in severe COVID disease

We next sought to determine whether CD4^+^ T cells produced high TNF-α in the lung in severe COVID-19 disease. To do this, we evaluated explanted lung tissue from a 51 y/o male who underwent bilateral lung transplantation for end-stage fibrotic lung disease following severe COVID-19 infection. We assessed lung mononuclear cells from bronchoalveolar lavage (BAL) fluid and lung parenchyma (LP) cells for CD4^+^/CD8^+^ ratios and S1-specific effector responses compared to PBMC from this patient. Further, we assessed BAL versus PBMC S1-specific responses on five lung transplant recipients with documented SARS-CoV-2 infection, all of which had severe COVID-19 respiratory disease (Table S2). As seen in PBMC from other severe COVID disease, the CD4^+^/CD8^+^ ratio was ≤1 in the PBMC, BAL and LP (Fig. 6A). Increased CD8^+^ T cell numbers are consistent with a prior report on BAL cells from severe COVID-19 patients(*23*). Compared to S1-specific responses from PBMC, CD4^+^ TNF-α > IFN-γ responses were significantly increased in the BAL and LP (Fig. 6B, C). Lung CD4^+^ T cells demonstrated CD69^+^CD103^+/−^CD45RA^-^CCR7^-^ resident memory phenotype, with increased CD38^hi^PD1^hi^Ki67^hi^ activation phenotype (Fig.6D, E). Together, these data support reduced CD4^+^ T cell numbers in the lung with severe COVID and increased S1-specific TNF-α production from activated CD4^+^ resident memory T cells.

**Figure 6.**
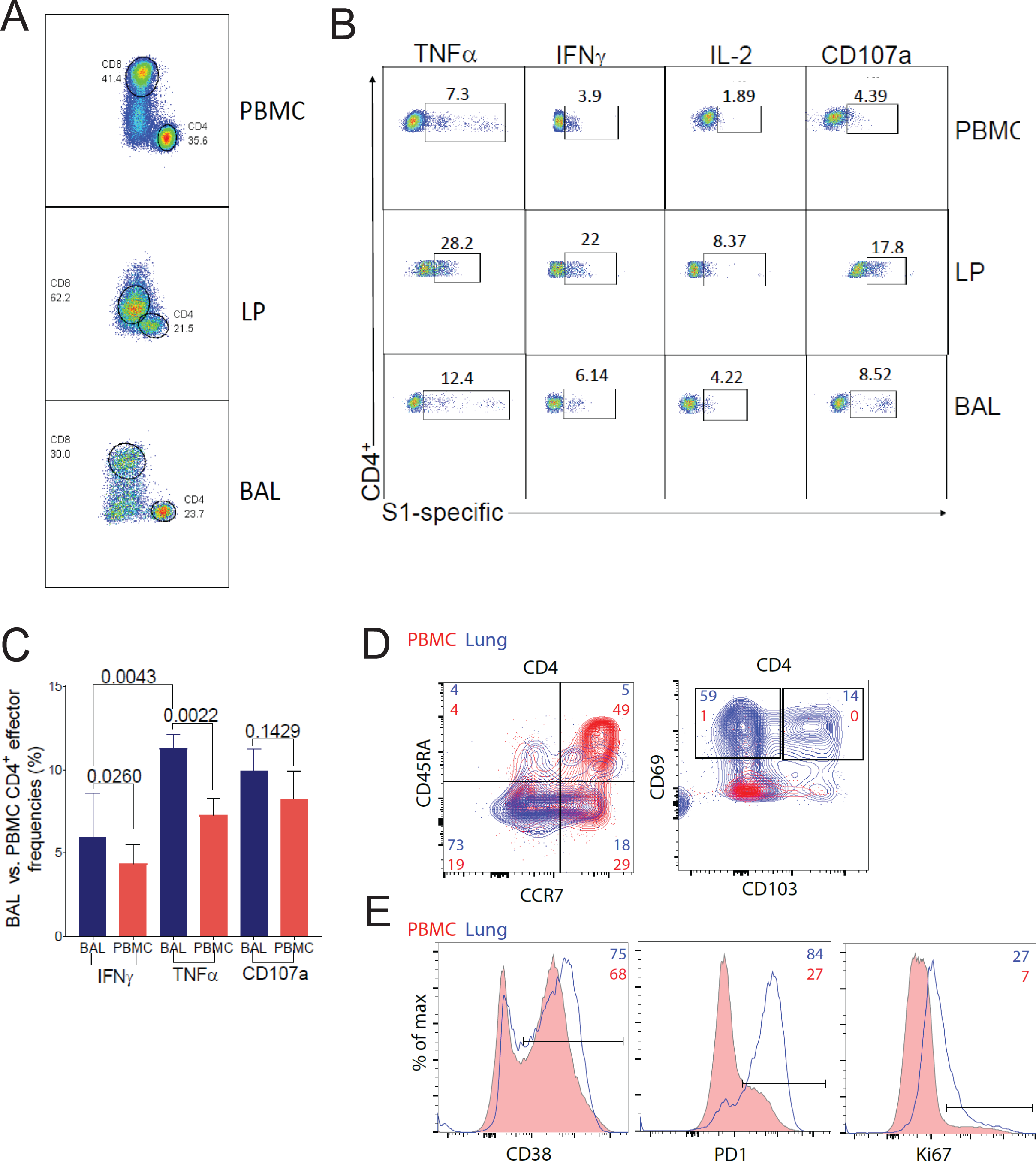
High TNF-α production by lung resident memory CD4^+^ T cells in severe COVID disease. *(A)*Representative flow cytometry plot on explanted lung tissue in severe COVID-19 disease. Flow cytometric representative plots of CD4^+^and CD8^+^ T cells in PBMC *(upper panel);* lung parenchyma (LP) *(middle panel)* and bronchoalveolar lavage (BAL) fluid-derived cells *(lower panel)* with cell frequencies of T cell subsets. *(B)* Representative flow cytometry plots of the same COVID-19 patient showing CD4^+^ T cells at 6 h after S1 peptides stimulation for TNFα, IFNγ, IL-2 and CD107a by intracellular cytokine staining in PBMC *(upper panels);* LP *(middle panels)* and BAL cells *(lower panels)* with cell frequencies shown. *(C)* Pooled data showing intracellular staining of 6 h S1-specific CD4^+^ T-cell effector responses IFNγ, TNFα, IL-2 and CD107a in BAL cells from n=6 as 5 lung transplant recipients and the explant above *(yellow columns)* vs. PBMC *(red columns)*. Bars represent median values ± interquartile range for S1-specific CD4^+^ frequencies and *p*-values were calculated using the Wilcoxon test with a two-sided p value of < 0.05 considered statistically significant.. *(E)* Representative flow cytometry plots of the COVID-19 lung explant showing CD4^+^ T cell suface expression of CD45RA^+^, CCR7^+^ *(left panel)* and CD69^+^CD103^+^ (*right panel)* on PBMC *(red)* or LP *(blue)* and values represent cell frequencies of T cell subsets. *(F)* Flow cytometry histograms of the COVID-19 explant showing LP CD4^+^ T cells CD38^+^ *(left panel)*, PD1^+^ *(middle panel)* and Ki67^+^ (*right panel)* overlayed LP (*blue lines*) on PBMC *(pink full lines)*.

### DISCUSSION

Herein, we show that severe COVID-19-associated lymphopenia is a predominant CD4^+^ lymphopenia and associated with an increased risk for mortality. Overall, severe COVID-19 disease was associated with a significant diminution in the peripheral CD4^+^/CD8^+^ ratios, whereas recovered, mild COVID-19, patients demonstrated a preservation of normal CD4^+^/CD8^+^ ratios. Based on these findings, we assessed the function of peripheral CD4^+^ T cells from severe COVID-19 versus mild recovered COVID-19 patients to evaluate immune mechanisms driving CD4^+^ lymphopenia. Indeed, we found that a disproportionate increase in TNF-α production and cytotoxic function from CD4^+^ T cells, but not CD8^+^ T cells, in response to the immunodominant antigen, S1, was evident in severe COVID-19 disease. We also found that S1-specific autocrine TNF-α-dependent production from CD4^+^ T cells themselves, and enhanced TNF-α responsiveness via TNFRI are key mechanisms leading to impaired CD4^+^ proliferation and activation induced cell death (AICD) and contribute to CD4^+^ lymphopenia. Together, our findings point to a skewed pro-inflammatory CD4^+^ T cell response among severe COVID-19 patients that is central to CD4^+^ T cell dysfunction and that plausibly contributes to the immunopathogenesis of severe disease.

While our studies found that S1 is the immunodominant antigen for SARS-CoV-2-induced T cell responses, we also detected significant effector T cell responses to other viral antigens, namely S2, NCAP and VEMP. However, CD4^+^ T cell responses to these antigens from severe disease patients did not demonstrate a hierarchical dominance of TNF-α to these antigens as the S1 response, whereas CD107a responses were similarly elevated to all antigens with severe disease. Further, S1 was also immunodominant for CD8^+^ T cell responses, however IFN-γ and CD107a were predominant in severe disease in contrast to CD4^+^ responses. Thus, S1-specific CD4^+^ T cell responses in severe disease were unique with increased levels of the pro-inflammatory effector cytokine TNF-α compared to mild disease controls, other antigens, and CD8^+^ T cells. Lastly, we show that CD4^+^TNF^+^ responses inversely correlated with absolute CD4^+^ T cell counts in severe disease. Together, these findings reveal that the immunodominant SARS-CoV-2 S1 protein elicits a strong TNF-α response from CD4^+^ T cells that subverts the host T cell response.

Earlier studies in HIV, measles and dengue viral infections have demonstrated that viremia results in impaired CD4^+^ T cell proliferation(*24–26*). While we cannot exclude SARS-CoV-2 replication having an impact on the reduced CD4^+^ proliferative responses observed in severe COVID-19 disease, we demonstrate that *in vitro* TNF-α blockade using infliximab, anti-TNFRI or other TNF-α antibodies strikingly restored these responses. Moreover, depleting CD8^+^ T cells and our finding of minimal spontaneous or S1-induced TNF-α from monocytes, supports the concept that autocrine CD4^+^ TNF-α production plays a major role in driving CD4^+^ dysfunction. Prior studies have shown mixed results on the impact of TNF-α on T cell responses. TNF-α can act as a co-stimulatory molecule and enhance T cell receptor (TCR)-dependent activation of T cells leading to increased cytokine and proliferative responses via NF-κB signaling(*27–30*). These co-stimulatory effects of TNF-α on T cell responses are predominantly mediated through TNFRII(*31, 32*). However, other studies have demonstrated *in vivo* that chronic exposure to TNF-α results in prolonged survival and attenuation of disease in NZB1/W F1 mice prone to lupus nephritis and suppresses type-I insulin-dependent diabetes mellitus in non-obese diabetic (NOD) mice(*33, 34*). Indeed, Cope et al. demonstrated that chronic TNF-α exposure inhibited T cell proliferation and cytokine production through TNFRI (p55 receptor) and that *in vitro* proliferative and Type-1 cytokine responses could be rescued using anti-TNF-α therapy, consistent with our observations(*35*). In contrast to our findings, other studies have found a T cell inhibitory role of TCR-dependent activation though the TNFRII and chronic TNF-α exposure(*36*). Additionally, chronic TNF-α exposure induced T cell hyporesponsiveness was subsequently shown to lead to impaired NF-κB signaling(*37*). Thus, differential effects of TNF-α on T cell responses and T cell-mediated pathology have been observed depending on the model system and duration of cytokine exposure. Our findings suggest that severe COVID-19 infection with persistent S1-specific TNF-α production from CD4^+^ T cells more closely models chronic TNF-α exposure, resulting in T cell hyporesponsiveness and dysfunction. In addition to anti-TNF-α blockade agents, we found that low dose exogenous IL-2 also rescued S1-specific proliferative responses in the setting of low IL-2 frequencies that significantly increased after 6 days in the presence of TNF-α blockade. Together, these data point to a relative IL-2-deficient state in severe COVID-19 disease that is TNF-α-dependent.

TNF-α can be a major regulatory pathway for apoptosis of various cells, as well as cell survival(*27, 38*). While the TNF-α/TNRI pathway has been demonstrated to be the predominant pathway for apoptosis in T cells, there is also evidence that the TNFRII plays a role(*39*). T cell apoptosis via the TNFRI and/or TNFRII has been shown to be important regulatory mechanism for the T cell response in viral infections such as lymphocytic choriomeningitis virus and influenza, as well as autoreactive and aged T cells(*40–43*). The TNF-α/TNFR pathways play highly complex direct and indirect roles in CD4^+^ and CD8^+^ T cell apoptosis during HIV infection, that include both TNFRI and TNFRII, in addition to other mechanisms(*44, 45*). Our studies show that TNF-α/TNFRI-dependent apoptosis of CD4^+^ T cells via AICD as a major mechanism contributing to CD4^+^ lymphopenia in severe SARS-CoV-2 infection. We also observed that TNFRI-mediated apoptosis in CD4^+^ T cells in severe COVID-19 was associated with increased surface expression of other activation markers such as CD95 (Fas) and CD38, and to a lesser extent PD-1, as previously reported in HIV infection(*46, 47*). We also observed that blockade of the Fas/FasL pathway, the major apoptotic pathway in HIV infection, also rescued CD4^+^ T cells from AICD, though to a lesser extent than TNF-α(*22, 48*). Our RNAseq studies also demonstrated that TNF-α blockade significantly reduced expression of Type-1 cytokines including TNF-α itself, NFκB signaling and FasL, consistent with the significant rescue of S1 re-stimulated CD4^+^ T cells from AICD. Taken together, our findings support high levels of TNF-α/TNFRI-dependent apoptosis of CD4^+^ T cells through AICD in severe COVID-19 disease that contributes to lymphopenia.

We evaluated lung resident T cells from an explanted lung (severe COVID-19) undergoing lung transplantation and found a similar diminution in the CD4^+^/CD8^+^ ratio in lung parenchymal and BAL cells, similar to PBMC. We further assessed BAL samples from four LTRs with recent severe COVID infection and had similar findings of reduced CD4^+^ frequencies in the lung, along with a similar hierarchy of increased CD4^+^ S1-specific TNF-α production, that was increased in both the LP and BAL compartments compared to the PBMC. These findings are consistent with our previous findings of enhanced CD4^+^ effector T cell responses in the lung during acute and chronic CMV infection, but unusual in its TNF-α predominance(*49*). Thus, our findings support a role for CD4^+^ T cell TNF-α production contributing to lung inflammation in severe COVID-19 disease and reduced CD4^+^ numbers in the lung.

Our studies found that the TNF-α blockade therapy, infliximab, a common therapy for rheumatoid arthritis and other autoimmune diseases, had a profound, dose response, *in vitro* effect in the restoration of S1-specific CD4^+^ proliferative responses and abrogation of S1-induced AICD(*50*). Indeed, recent studies have shown that treatment with the IL-6R antagonist tocilizumab resulted in increased circulating lymphocytes and cytotoxic NK cells(*19, 51*). Moreover, high IL-6 and TNF-α levels on hospital admission correlated with mortality in severe COVID-19 disease(*52*). Some observational patient data suggest a potential survival benefit from poor COVID-19 outcomes in patients already receiving TNF-α blockade therapy compared to patients on other immunomodulators and a mouse SARS-CoV-2 infection model demonstrates improved survival with TNF-α blockade(*53*). A current NIH Phase 3 trial is evaluating infliximab along with other immunomodulators (ACTIV-1 trial) for moderate to severe COVID-19 disease; https://www.nih.gov/research-training/medical-research-initiatives/activ/covid-19-therapeutics-prioritized-testing-clinical-trials. Together, our studies and other lines of evidence suggest a potential role for TNF-α blockade therapy in severe COVID-19 respiratory disease.

In summary, we show that the CD4^+^ T cell immunodominant response to S1 during severe COVID induces high TNF-α levels resulting in autocrine TNF-α/TNFRI-dependent impaired CD4^+^ proliferation and susceptibility to AICD. We further found that high autologous CD4^+^ TNF-α production correlated with a predominant CD4^+^ lymphopenia during severe COVID disease, which we show is associated with increased mortality. Importantly, lung resident T cells produce elevated TNF-α along with reduced CD4^+^ T cell numbers. Together, our studies demonstrate mechanisms leading to COVID-associated lymphopenia and provide a strong rationale for testing TNF-α blockade therapy in severe COVID-19 disease.

### MATERIAL AND METHODS

#### Study participants

Patients from the University of Pittsburgh Medical Center hospitals admitted with COVID-19 or screened for convalescent plasma donation were identified and provided informed written consent for participation in a UPMC Institutional Review Board-approved protocol, at the University of Pittsburgh.

#### Preparation of PBMC, BAL and lung parenchyma cells

Blood samples were obtained from outpatient and inpatient study subjects as above whenever possible and PBMC isolated from heparinized blood samples by density gradient centrifugation using Ficoll-Paque (Cytiva, Sweden), aliquoted and stored in freezing medium in liquid nitrogen. BAL cells were isolated from excess BAL fluid from n=6 patients (5 lung transplant recipients and 1 severe COVID-19 lung explant) by centrifugation. Lung parenchymal mononuclear cells were purified as previously described(*54*). Lung parenchymal and BAL cells were aliquoted and stored in freezing medium in liquid nitrogen.

#### Flow cytometric studies

##### Surface and intracellular cytokine staining

*In vitro* 6 h stimulation and intracellular cytokine staining (ICS) for IFN-γ, TNF-α, IL-2, IL-17a, IL-13 and CD107a expression was determined using pools of overlapping 15-mer peptides of SARS-CoV-2 specific for S1, S2, VEMP and NCAP (JPT Peptide Technologies Inc., Germany). SEB or pooled COVID-19 specific peptides were used at (1μg/mL) using 10^6^ cells per condition for 6 h at 37°C, 5% CO_2_. All re-stimulations for ICS were performed using 10^6^ cells with brefeldin-A (10 μg/mL) and monensin (5 μg/ml) with anti-CD107a-Pac-Blue added at the beginning of culture. Surface staining was performed using the fluorochrome-labeled antibodies: anti-CD3-Alexa-Fluor700 (UCHT1), CD14-BV605 (MSE2), anti-CD8-V500 (SK1), anti-CD4-APC-Cy7 (SK3) (BD Biosciences, San Jose, CA), Live/Dead Fixable Blue Dead-Cell Stain (Invitrogen, Thermo Fisher Scientific, Waltham, MA), TNFRI (CD120a)-PE (clone W15099A), TNFRII (CD120b)-APC (clone 3G7A02), CD95-FITC (clone DX2), PD1-PE or BV450 (clone EH1.2.1), CD38-APC (clone HB-7) (BD-Biosciences San Jose, CA and BioLegend, San Diego, CA). For Lung cell experiments, the following antibodies were used: CD3 -BUV395, CD4-BV480, CD8-BUV737 (BD Biosciences, San Jose, CA), CD45RA-BV605, CCR7-BV785, CD69-BV650, PD1-BV711, CD103-APC-Cy7, CD38-PED594 and Ki67-AF499 (BioLegend, San Diego, CA). ICS was performed using anti-IFN-γ-BV605, anti-TNF-α-PE-Cy7, anti-IL-2-BV650, anti-CD107a-PECF594 or BV450 (clone H4A3), anti-IL-13-BV421 (clone JES10-5A2) (BD-Biosciences, San Jose, CA and BioLegend, San Diego, CA) and anti-IL-17a-APC (clone eBio64Dec17) (eBioscience, Termo Fisher Scientific, Waltham, MA). Using Boolean gating analysis, cytokine co-expression was determined using *SPICE* software (version 6.1), downloaded from http://exon.niaid.nih.gov/spice.

##### *In vitro* antigen-specific T cell proliferation

Thawed frozen PBMCs were labeled with 0.2 mM CFSE (Invitrogen, Carlsbad, CA) in PBS for 7 min. and blocked with FCS for 2 min. The cells were immediately washed and plated in complete RPMI supplemented with 10% FCS in a 96 deep well plate for 6 days. For antigen-specific stimulation, PBMCs were stimulated in the presence or absence of SARS-CoV-2 peptide pools mix (S1, S2, VEMP and NCAP; JPT, Berlin, Germany) at (1μg/mL) for 6 days. Isolated cells underwent secondary re-stimulation *in vitro* for 6 h with/without S1 peptide pool and assessed by flow cytometry for CFSE dilution and ICS. All gates for cytokine frequencies were set using the medium alone control and subtracted from peptide-stimulated sample frequencies. All cells were collected for flow cytometric analysis using an LSR Fortessa-cytometer with UV laser (BD Biosciences, Franklin Lakes, NJ) or Cytek Aurora Flow-cytomer (Cytek, Bethesda, MD). Data analysis and graphic representations were done with *FlowJo* v.10 7.2 (BD-Becton Dickinson & Company, Ashland, OR).

#### Bead depletion of CD8^+^ T cells and negative selection for CD4^+^ T cell enrichment in PBMC

In select experiments (proliferation and apoptosis studies), we performed CD8^+^ depletion using the human kit II by EasySep™ human CD8 Kit II Cell Separation (17853-STEM Cell Technologies, Cambridge, MA) to enrich PBMC for CD4^+^ T cells. For single cell RNA sequencing experiments, we used peripheral CD4^+^ T cells purify by negative selection using the kit EasySep™ Human CD4^+^ T Cell Enrichment Kit Immunomagnetic negative selection (17952-STEM Cell Technologies, Cambridge, MA) to enrich PBMC for CD4^+^ T cells.

#### Antibody blocking studies

In select experiments we used anti-human TNF-α and isotype IgG1 (Pharmingen, BD-Biosciences, San Jose, CA), Infliximab (anti-human TNF-α chimeric monoclonal antibody; Janssen, Horsham, PA), anti-TNFRI (clone 16803), anti-TNFRII (clone 22210) **(R&D Systems, Inc.,** Minneapolis, MN), and anti-Fas (clone ZB4) (Millipore Sigma, Burlington, MA).

#### Apoptosis studies

In apoptosis studies, cells were stained with anti-Annexin V-Brilliant Violet-450 after in vitro stimulation with S1 peptides with/without various blocking or isotype control (IgG1) Abs, for 12 hours at 37°C. In select experiments, anti-Fas IgG1 (clone-ZB4; neutralizing) at 10 μg/ml plate-bound for 12 hours at 37°C or isotype control Abs IgG1 (Millipore). All cells were collected for flow cytometric analysis using an LSR Fortessa-cytometer with UV laser (BD Biosciences, San Jose, CA). Data analysis and graphic representations were done with FlowJo v.10 7.2 ((BD-Becton Dickinson & Company, Ashland, OR).

#### Single cell RNA analysis

We used purified CD4^+^ T cells by negative selection (17952-STEM Cell Technologies, Cambridge, MA) to enrich PBMC for CD4^+^ T cells 3 h following S1-stimulation in the presence or absence of infliximab. For the RNAseq analysis, differential gene expression analysis between subjects was completed using CLC Genomics Workbench version 20. Enriched CD4^+^ T cells were divided into four samples for HTO labeling. Each sample was subjected to the Totalseq-C and cell-hashing protocol, targeting a recovery of ∼5000 cells per sample(*55*). All HTO-tagged cells were pooled together and prepared using the 10X Genomics with 5’ Gel Bead Kit V1. The prepared library was subsequently sequenced on Illumina Novaseq platform with a depth of 50K read per cell. Downstream analysis of single cell RNA was performed using Seurat v3(*56*) and R software(*57*), including permissive filtering of low-quality cells, normalization, identification of highly variable genes, and Louvain clustering. Cell-cluster identity was established using previously reported immune gene signatures(*7*). Subsets of singlets were isolated based on upregulation of *CD3E* (expression threshold >0.5), and downregulation of *CD68* (<0.5), *CD19* (<0.5), and *CD8A* (<0.5). The remaining cells were treated as CD4^+^ T cells. Normalization and Louvain clustering was then performed on this purified subset of singlets. Highly variable genes between clusters of CD4^+^ T cells were identified based on the non-parametric Wilcoxon rank sum test. Top differentially expressed genes across clusters were visualized using the DoHeatmap function in Seurat. Feature plots were performed on key genes identified. Gene set enrichment analysis was performed using Fast Gene Set Enrichment Analysis (fgsea) https://www.biorxiv.org/content/10.1101/060012v3.full differentially expressed genes between clusters 1 and 3 among the CD4^+^ T cell UMAP were included as input. Genes were references to the Hallmark pathways from MSigDB(*58*). Pathways were included if the absolute value of the normalized enrichment score > 1.5.

#### Statistical analysis

Distributions of all measured variables were performed using nonparametric testing as indicated. The non-parametric tests of Wilcoxon signed-rank and Mann-Whitney U analysis were applied using GraphPad Prism 9 (La Jolla, CA). A two-tailed P < 0.05 was considered statistically significant. All correlations were determined using Spearman rho test and Spearman rank correlation test.

## Acknowledgements

We wish to thank all of the UPMC physicians, nurses and other hospital staff for their tireless, hard work and dedication to the care of COVID-19 patients during this pandemic.

## Funding

The Pittsburgh Foundation, The David Scaife Foundation, NIH R01 HL133184 (JM).

## Author contributions

J.F.M. obtained funding and contributed study design and writing of manuscript

I.P., M.E.S., K.C. contributed to study design, performed experiments and writing of manuscript

J.K.A., B.B.C. contributed to data analysis and manuscript preparation

A.D., M.J.B., E.J.L., S.C.L., J.C.S., X.A. assisted I.P., M.E.S. and K.C. in performing immunologic experiments

C.J.I., X.C., G.K., I.K., M.S., D.J.K. contributed to clinical data extraction and analysis and manuscript preparation

R.K., R.B., S.J.H., K.L., A.M., B.J.M., J.E.K., A.H.W., D.K.M., D.J.T. contributed to patient screening, identification and study enrollment and clinical data collection

E.A.L., S.K., B.J., M.R.M., J.M.P., P.G.S. contributed to lung transplant patient identification and study sample collections

## Competing interests

The authors declare no competing interests.

## Data and materials availability

All data are available in the main text or the supplementary materials.

---

**TABLE S1.**

|  | **ALC < 700**<br>**(n = 70)** | **ALC ≥ 700**<br>**(n = 78)** | **p-value** |
| --- | --- | --- | --- |
| **Age, median [IQR]** | 71.0 (60-78) | 66.0 (54-76) | 0.07 |
| **Race** |  |  | 0.40 |
| White | 56 (80.0%) | 68 (87.2%) |  |
| Black | 9 ( 2.8%) | 8 (10.3%) |  |
| Other | 1 (1.4%) | 1 (1.3%) |  |
| Unknown/No Answer | 4 (5.8%) | 1 (1.3%) |  |
| **Female** | 26 (37.1%) | 31 (39.7%) | 0.74 |
| **Peak Resp Support** |  |  | **0.02** |
| None | 1 (1.4%) | 7 (9.0%) |  |
| Low flow NC | 15 (21.4%) | 21 (26.9%) |  |
| High low NC | 21 (30.0%) | 24 (30.8%) |  |
| NIPPV | 3 (4.3%) | 4 (5.1%) |  |
| Mechanical<br>Ventilation | 30 (42.9%) | 20 (25.6%) |  |
| ECMO | 0 (0%) | 1 (1.2%) |  |
| **Cardiovascular Disease** | 48 (68.6%) | 43 (55.1%) | 0.09 |
| **Respiratory Disease** | 19 (27.1%) | 17 (21.8%) | 0.45 |
| **Diabetes** | 27 (38.6%) | 30 (38.5%) | 0.99 |

ALC - absolute lymphocyte count; NC – nasal cannula; NIPPV – non-invasive partial pressure ventilation; ECMO – extracorporeal membrane oxygenation

**TABLE S2.**
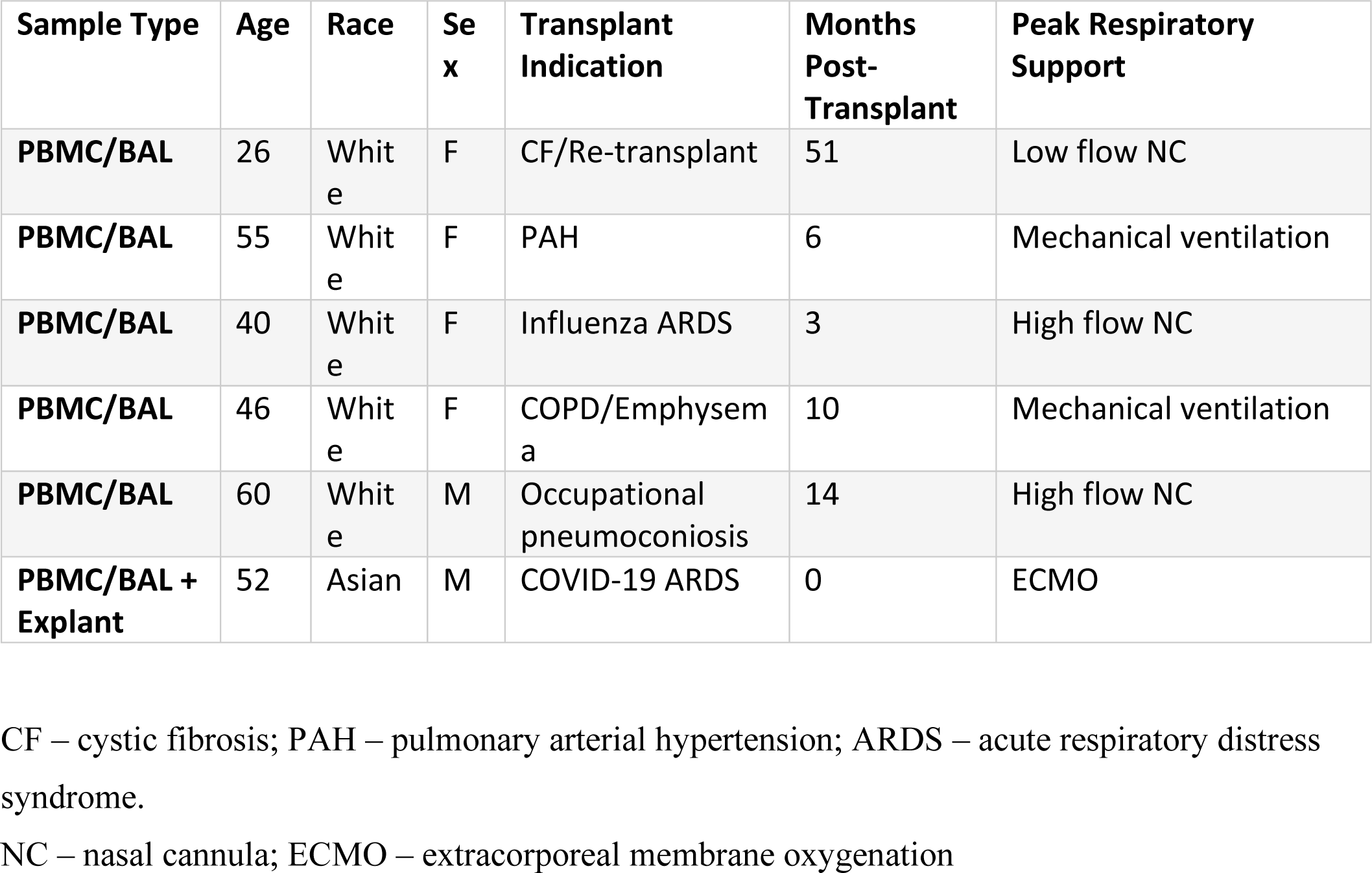

| **Sample Type** | **Age** | **Race** | **Sex** | **Transplant Indication** | **Months Post-Transplant** | **Peak Respiratory Support** |
| --- | --- | --- | --- | --- | --- | --- |
| **PBMC/BAL** | 26 | White | F | CF/Re-transplant | 51 | Low flow NC |
| **PBMC/BAL** | 55 | White | F | PAH | 6 | Mechanical ventilation |
| **PBMC/BAL** | 40 | White | F | Influenza ARDS | 3 | High flow NC |
| **PBMC/BAL** | 46 | White | F | COPD/Emphysema | 10 | Mechanical ventilation |
| **PBMC/BAL** | 60 | White | M | Occupational pneumoconiosis | 14 | High flow NC |
| **PBMC/BAL + Explant** | 52 | Asian | M | COVID-19 ARDS | 0 | ECMO |

CF – cystic fibrosis; PAH – pulmonary arterial hypertension; ARDS – acute respiratory distress syndrome.
NC – nasal cannula; ECMO – extracorporeal membrane oxygenation

## Supplementary Materials

**Figure S1.**
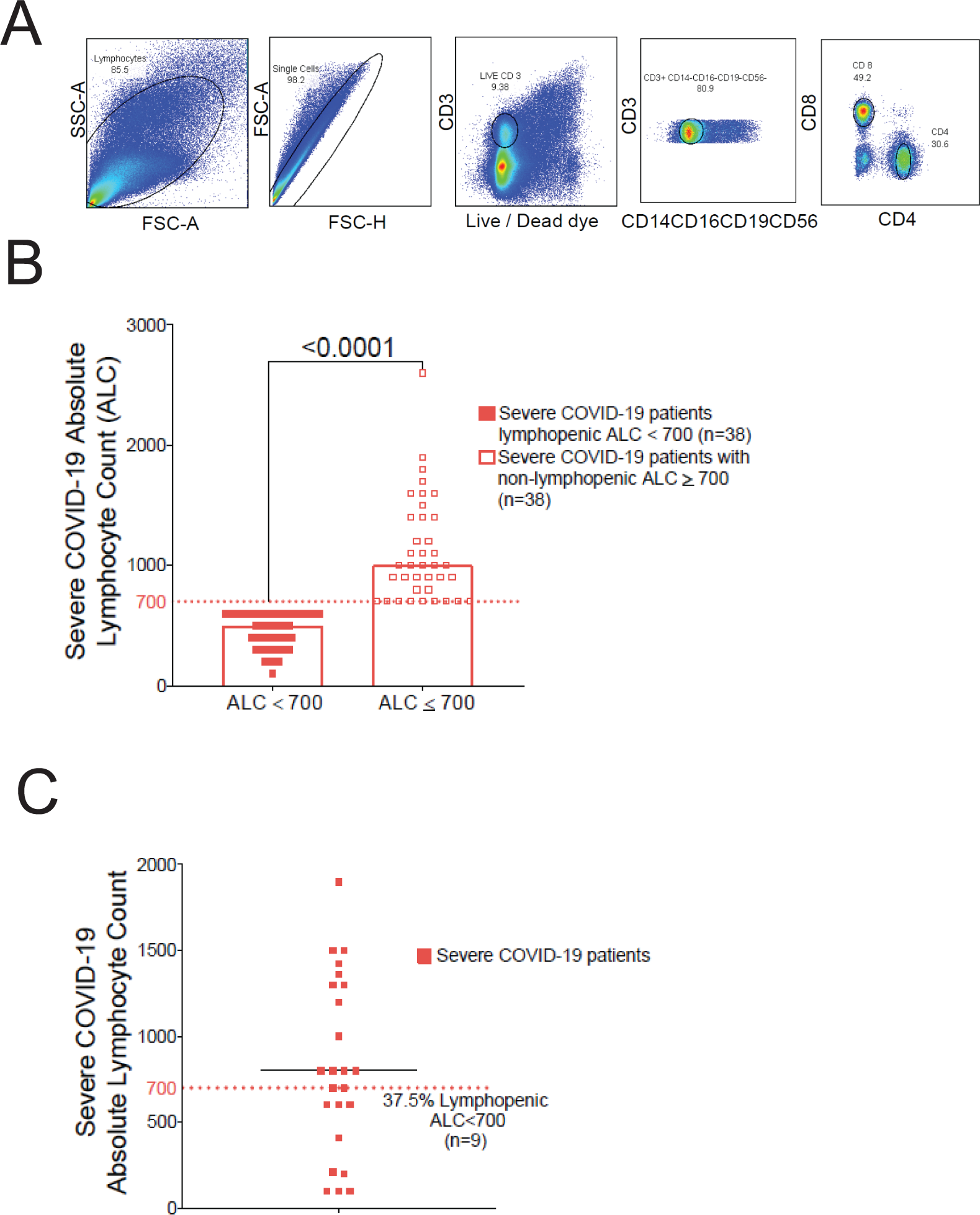
Flow Cytometric Gating and Absolute Lymphocyte Counts. *(A)* Representative flow plots showing the gating strategy used in the flow cytometric analysis (FlowJo software). *(B)* Pooled data (n=76) showing absolute lymphocyte count (ALC) data from severe COVID-19 with lymphopenia (ALC<700) (n=38) *(full red squares)* compared with non-lymphopenia (ALC≥ 700) patients (n=38) *(empty red squares)*. *(C)* Cumulative absolute lymphocyte count (ALC) data from the cohort used in functional studies of severe COVID-19 (n=24) *(red squares)*.

**Figure S2.**
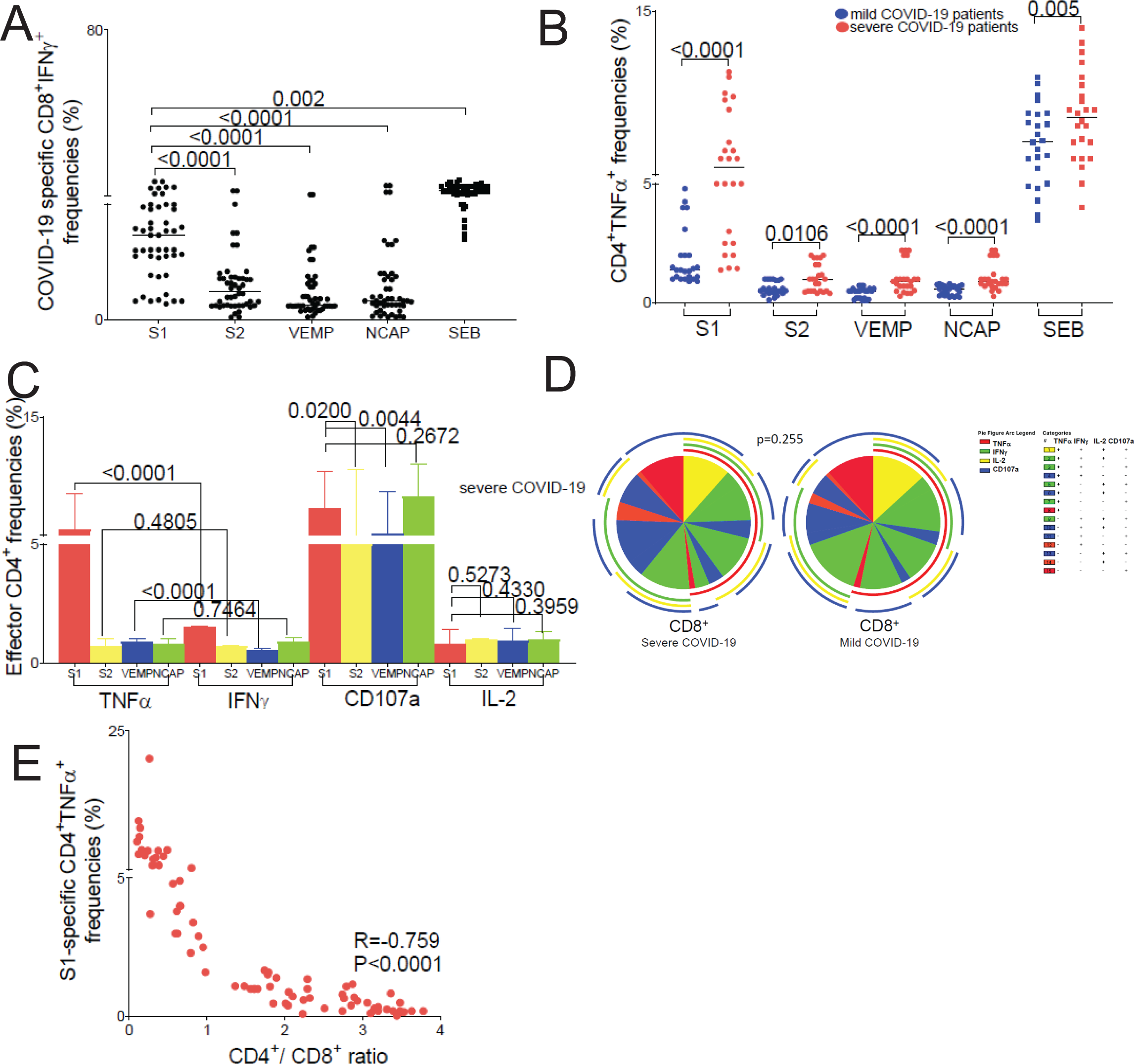
Supplemental CD4^+^ and CD8^+^ effector cytokine responses. *(A)* Pooled data of the CD8^+^ T-cell IFNγ^+^ immunodominance in COVID-19 cohort (n=48) for S1, S2, VEMP and NCAP-specific and SEB-reactive effector responses. *(B)* Cumulative data of the CD4^+^ T-cell TNFα^+^ in mild *(blue dots)* and severe *(red dots)* COVID-19 cohort (n=48) for S1, S2, VEMP and NCAP - specific and SEB-reactive effector responses. *(C)* Pooled data of the CD4^+^ T-cell COVID-19 antigen specific [S1 *(red columns)*, S2 *(yellow columns)*, VEMP *(blue columns)* and NCAP *(green columns)*] cytokines (TNFα, IFNγ, CD107a and IL-2) responses in severe COVID-19 cohort (n=24). Bars represent median values, and *P* values were calculated using the Mann-Whitney-test, where p value of < 0.05 considered statistically significant. *(D)* Individual pie charts showing CD8^+^ on severe COVID-19 disease *(left pie)* and CD8^+^ for mild COVID-19 disease *(right pie)* T-cell multifunctional responses (n=48). The CD8^+^ multifunctional responses are for four cytokines: TNFα (*red arch*), IFNγ (*green arch*), IL-2 (*yellow arch*) and CD107a (*blue arch*) and their combination of cytokine responses: +4-*yellow pie fraction*; +3-*green pie fraction;* +2-*blue pie fraction* and +1-*red pie fraction*. Using Boolean analysis, the percentage of total and individual effector multifunctional subset responses. Significant differences when comparing mean frequencies of single and multifunctional responses are indicated by a p *≤* 0.05. All p-values were determined by the Kruskal-Wallis one-way ANOVA or Wilcoxon signed-rank test. Data analyzed using the program *SPICE*. *(E)* Inverse correlation of S1-specific COVID-19 CD4^+^TNFα^+^ response frequencies with CD4^+^/CD8^+^ ratio on n=76 severe COVID-19 *(red dots).* Analysis was performed using the Spearman rho test and correlation coefficient (R) and (P) values were calculated using Spearman rank correlation test.

**Figure S3.**
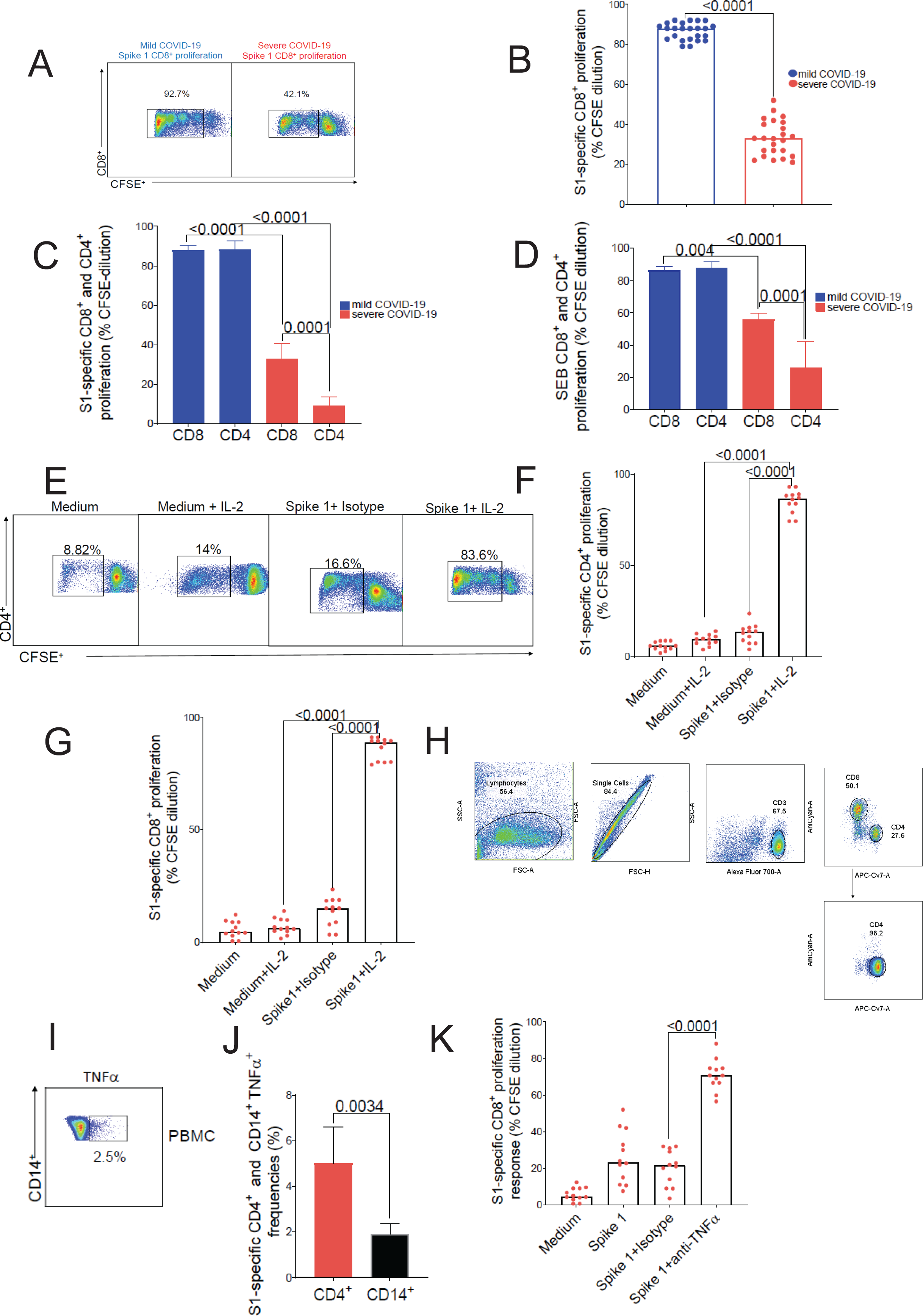
Supplemental T cell Proliferation, Cytokine and CD8^+^ depletion data. Representative flow cytometric plots *(A)* and cumulative data *(B)* showing S1-specific CD8^+^ T cell proliferation by CFSE dilution in mild *(left panel-A and blue dots-B)* vs. severe *(right panel-A and red dots-B)* COVID-19. The numbers indicate frequencies of live-gated CFSE diluted events (%). Bars represent median values and *p*-values were calculated using the Mann-Whitney-Wilcoxon test. *(C)* Cumulative data showing S1-specific CD8^+^ vs. CD4^+^T cell proliferation by CFSE dilution in mild *(blue columns)* vs. severe *(red columns)* COVID-19. Bars represent median values and *p*-values were calculated using the Mann-Whitney-Wilcoxon test. *(D)* Cumulative data showing SEB-reactive CD8^+^ vs. CD4^+^T cell proliferation by CFSE dilution in mild *(blue columns)* vs. severe *(red columns)* COVID-19. Bars represent median values and *p*-values were calculated using the Mann-Whitney-Wilcoxon test. Addition of exogenous IL-2 (20 IU/ml) to PBMC cultures and day-6 S1-specific CD4^+^ T cell proliferation with representative flow cytometric plots *(E)* and cumulative data *(F)* showing S1-specific CD4^+^ T cell proliferation by CFSE dilution in severe disease versus controls, (n=12) *(red dots). C*umulative data showing S1-specific CD8^+^ T cell proliferation by CFSE dilution using exogenous IL-2 *(G)* in severe COVID-19 patients, (n=12) *(red dots).* Bars represent median values and *p*-values were calculated using the Mann-Whitney-Wilcoxon test. *(H)* Representative gating strategy and analysis of the CD8^+^-depleted PBMC used in some proliferation and apoptosis experiments. Representative flow cytometric plots *(I)* and cumulative data *(J)* showing S1-specific CD14^+^ TNFα^+^ on from severe COVID-19 patients, (n=12). The numbers indicate frequencies of TNFα^+^events (%) and bars represent median values of S1-specific CD4^+^TNFα^+^ *(red columns)* and CD14^+^TNFα^+^ and *p*-values were calculated using the Mann-Whitney-Wilcoxon test.

**Figure S4.**
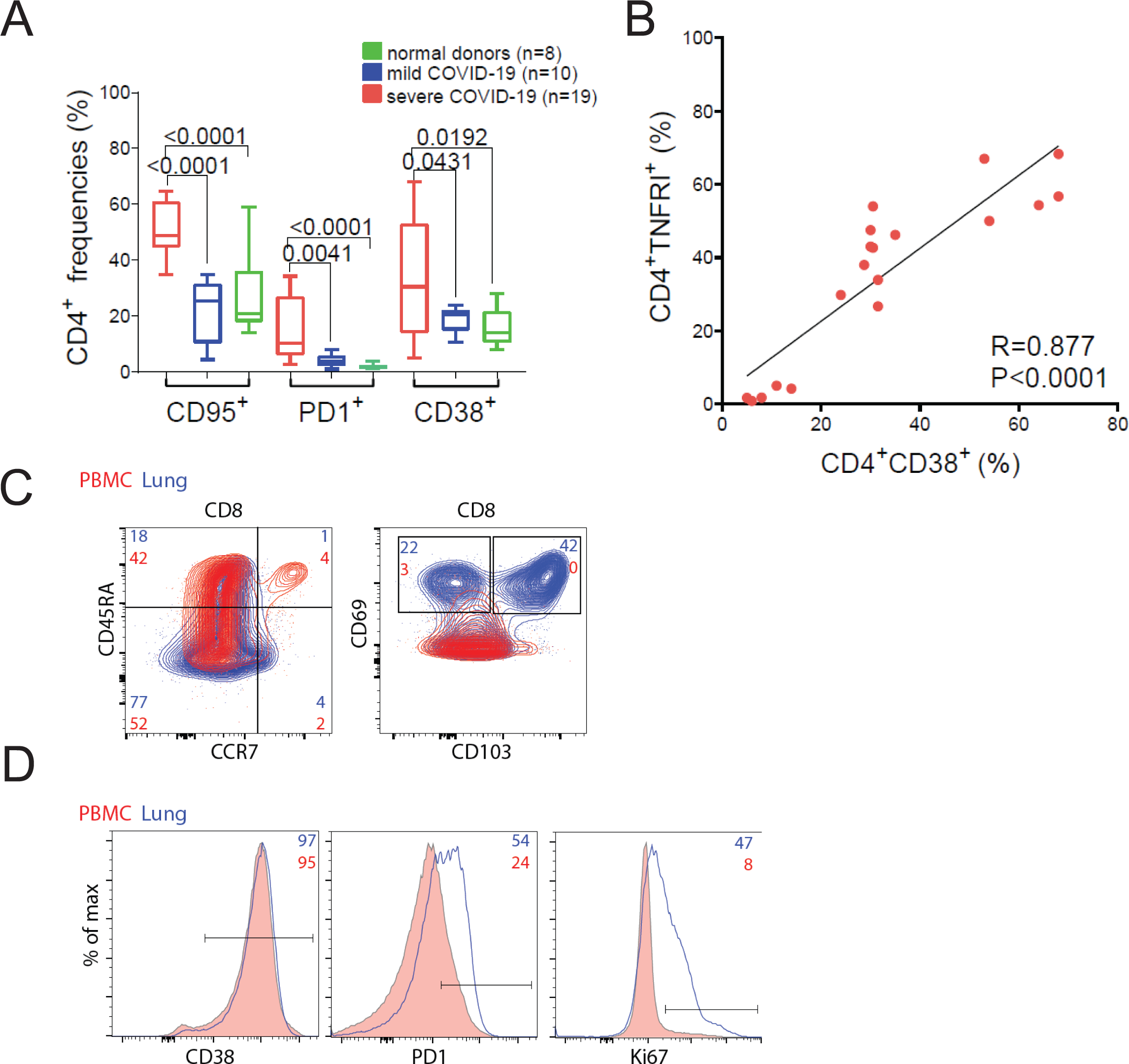
Supplemental PBMC and Lung T cell phenotyping. *(A)*Cumulative data showing CD4^+^ CD95^+^ (Fas), PD1^+^ and CD38^+^ surface markers expression in severe *(red columns)* n=19 vs. mild *(blue columns)* COVID-19 n=10 vs. normal donors *(green columns)* n=8*. (B)* Correlation of CD4^+^CD38^+^ surface expression with TNFRI^+^ in severe COVID-19 patients (n=19). Analysis was performed using the Spearman rho test and correlation coefficient (R) and (P) values were calculated using Spearman rank correlation test. *(C)* Representative flow cytometry plots of the COVID-19 lung explant showing CD8^+^ T cell surface expression of CD45RA^+^, CCR7^+^ *(left panel)* and CD69^+^CD103^+^ (*right panel)* on PBMC *(red)* or Lung Parenchima (LP) *(blue)* and values represent cell frequencies of T cell subsets. *(D)* Flow cytometry histograms of the COVID-19 explant showing LP CD8^+^ T cells CD38^+^ *(left panel)*, PD1^+^ *(middle panel)* and Ki67^+^ (*right panel)* overlayed LP (*blue lines*) on PBMC *(pink full lines)*.

## Notes

### Competing Interest Statement

The authors have declared no competing interest.

## References

1. R. H. Du et al., Predictors of mortality for patients with COVID-19 pneumonia caused by SARS-CoV-2: a prospective cohort study. Eur Respir J 55, (2020).

2. H. J. Yang, Y. M. Zhang, M. Yang, X. Huang, Predictors of mortality for patients with COVID-19 pneumonia caused by SARS-CoV-2. Eur Respir J 56, (2020).

3. M. J. Cummings et al., Epidemiology, clinical course, and outcomes of critically ill adults with COVID-19 in New York City: a prospective cohort study. medRxiv, (2020).

4. D. C. Fajgenbaum, C. H. June, Cytokine Storm. N Engl J Med 383, 2255–2273 (2020).

5. P. Sinha, M. A. Matthay, C. S. Calfee, Is a “Cytokine Storm” Relevant to COVID-19? JAMA Intern Med 180, 1152–1154 (2020).

6. R. C. Group et al., Dexamethasone in Hospitalized Patients with Covid-19. N Engl J Med 384, 693–704 (2021).

7. D. C. Angus et al., Effect of Hydrocortisone on Mortality and Organ Support in Patients With Severe COVID-19: The REMAP-CAP COVID-19 Corticosteroid Domain Randomized Clinical Trial. JAMA 324, 1317–1329 (2020).

8. R.-C. Investigators et al., Interleukin-6 Receptor Antagonists in Critically Ill Patients with Covid-19. N Engl J Med, (2021).

9. C. Salama et al., Tocilizumab in Patients Hospitalized with Covid-19 Pneumonia. N Engl J Med 384, 20–30 (2021).

10. S. Gupta et al., Association Between Early Treatment With Tocilizumab and Mortality Among Critically Ill Patients With COVID-19. JAMA Intern Med 181, 41–51 (2021).

11. V. C. Veiga et al., Effect of tocilizumab on clinical outcomes at 15 days in patients with severe or critical coronavirus disease 2019: randomised controlled trial. BMJ 372, n84 (2021).

12. J. H. Stone et al., Efficacy of Tocilizumab in Patients Hospitalized with Covid-19. N Engl J Med 383, 2333–2344 (2020).

13. I. O. Rosas et al., Tocilizumab in Hospitalized Patients with Severe Covid-19 Pneumonia. N Engl J Med, (2021).

14. L. Tan et al., Lymphopenia predicts disease severity of COVID-19: a descriptive and predictive study. Signal Transduct Target Ther 5, 33 (2020).

15. Z. Chen, E. John Wherry, T cell responses in patients with COVID-19. Nat Rev Immunol 20, 529–536 (2020).

16. D. Mathew et al., Deep immune profiling of COVID-19 patients reveals distinct immunotypes with therapeutic implications. Science 369, (2020).

17. L. Kuri-Cervantes et al., Comprehensive mapping of immune perturbations associated with severe COVID-19. Sci Immunol 5, (2020).

18. B. Diao et al., Reduction and Functional Exhaustion of T Cells in Patients With Coronavirus Disease 2019 (COVID-19). Front Immunol 11, 827 (2020).

19. A. Mazzoni et al., Impaired immune cell cytotoxicity in severe COVID-19 is IL-6 dependent. J Clin Invest 130, 4694–4703 (2020).

20. A. Grifoni et al., Targets of T Cell Responses to SARS-CoV-2 Coronavirus in Humans with COVID-19 Disease and Unexposed Individuals. Cell 181, 1489–1501 e1415 (2020).

21. A. Tarke et al., Comprehensive analysis of T cell immunodominance and immunoprevalence of SARS-CoV-2 epitopes in COVID-19 cases. bioRxiv, (2020).

22. I. Popescu et al., Activation-induced cell death drives profound lung CD4(+) T-cell depletion in HIV-associated chronic obstructive pulmonary disease. Am J Respir Crit Care Med 190, 744–755 (2014).

23. M. Liao et al., Single-cell landscape of bronchoalveolar immune cells in patients with COVID-19. Nat Med 26, 842–844 (2020).

24. A. C. McNeil et al., High-level HIV-1 viremia suppresses viral antigen-specific CD4(+) T cell proliferation. Proc Natl Acad Sci U S A 98, 13878–13883 (2001).

25. H. C. Whittle, J. Dossetor, A. Oduloju, A. D. Bryceson, B. M. Greenwood, Cell-mediated immunity during natural measles infection. J Clin Invest 62, 678–684 (1978).

26. A. Mathew et al., Impaired T cell proliferation in acute dengue infection. J Immunol 162, 5609–5615 (1999).

27. A. K. Mehta, D. T. Gracias, M. Croft, TNF activity and T cells. Cytokine 101, 14–18 (2018).

28. P. Scheurich, B. Thoma, U. Ucer, K. Pfizenmaier, Immunoregulatory activity of recombinant human tumor necrosis factor (TNF)-alpha: induction of TNF receptors on human T cells and TNF-alpha-mediated enhancement of T cell responses. J Immunol 138, 1786–1790 (1987).

29. S. Yokota, T. D. Geppert, P. E. Lipsky, Enhancement of antigen- and mitogen-induced human T lymphocyte proliferation by tumor necrosis factor-alpha. J Immunol 140, 531–536 (1988).

30. D. Banerjee, H. C. Liou, R. Sen, c-Rel-dependent priming of naive T cells by inflammatory cytokines. Immunity 23, 445–458 (2005).

31. L. A. Tartaglia et al., Stimulation of human T-cell proliferation by specific activation of the 75-kDa tumor necrosis factor receptor. J Immunol 151, 4637–4641 (1993).

32. R. M. Aspalter, M. M. Eibl, H. M. Wolf, Regulation of TCR-mediated T cell activation by TNF-RII. J Leukoc Biol 74, 572–582 (2003).

33. C. O. Jacob, H. O. McDevitt, Tumour necrosis factor-alpha in murine autoimmune ‘lupus’ nephritis. Nature 331, 356–358 (1988).

34. C. O. Jacob, S. Aiso, S. A. Michie, H. O. McDevitt, H. Acha-Orbea, Prevention of diabetes in nonobese diabetic mice by tumor necrosis factor (TNF): similarities between TNF-alpha and interleukin 1. Proc Natl Acad Sci U S A 87, 968–972 (1990).

35. A. P. Cope et al., Chronic tumor necrosis factor alters T cell responses by attenuating T cell receptor signaling. J Exp Med 185, 1573–1584 (1997).

36. R. M. Aspalter, H. M. Wolf, M. M. Eibl, Chronic TNF-alpha exposure impairs TCR-signaling via TNF-RII but not TNF-RI. Cell Immunol 237, 55–67 (2005).

37. L. F. Lee et al., Genomic expression profiling of TNF-alpha-treated BDC2.5 diabetogenic CD4+ T cells. Proc Natl Acad Sci U S A 105, 10107–10112 (2008).

38. D. Brenner, H. Blaser, T. W. Mak, Regulation of tumour necrosis factor signalling: live or let die. Nat Rev Immunol 15, 362–374 (2015).

39. X. Li, Y. Yang, J. D. Ashwell, TNF-RII and c-IAP1 mediate ubiquitination and degradation of TRAF2. Nature 416, 345–347 (2002).

40. M. Suresh, A. Singh, C. Fischer, Role of tumor necrosis factor receptors in regulating CD8 T-cell responses during acute lymphocytic choriomeningitis virus infection. J Virol 79, 202–213 (2005).

41. M. E. Wortzman, G. H. Lin, T. H. Watts, Intrinsic TNF/TNFR2 interactions fine-tune the CD8 T cell response to respiratory influenza virus infection in mice. PLoS One 8, e68911 (2013).

42. L. Ban et al., Selective death of autoreactive T cells in human diabetes by TNF or TNF receptor 2 agonism. Proc Natl Acad Sci U S A 105, 13644–13649 (2008).

43. S. Aggarwal, S. Gollapudi, S. Gupta, Increased TNF-alpha-induced apoptosis in lymphocytes from aged humans: changes in TNF-alpha receptor expression and activation of caspases. J Immunol 162, 2154–2161 (1999).

44. L. M. de Oliveira Pinto, S. Garcia, H. Lecoeur, C. Rapp, M. L. Gougeon, Increased sensitivity of T lymphocytes to tumor necrosis factor receptor 1 (TNFR1)- and TNFR2-mediated apoptosis in HIV infection: relation to expression of Bcl-2 and active caspase-8 and caspase-3. Blood 99, 1666–1675 (2002).

45. G. Herbein, K. A. Khan, Is HIV infection a TNF receptor signalling-driven disease? Trends Immunol 29, 61–67 (2008).

46. C. A. Muro-Cacho, G. Pantaleo, A. S. Fauci, Analysis of apoptosis in lymph nodes of HIV-infected persons. Intensity of apoptosis correlates with the general state of activation of the lymphoid tissue and not with stage of disease or viral burden. J Immunol 154, 5555–5566 (1995).

47. M. L. Gougeon et al., Programmed cell death in peripheral lymphocytes from HIV-infected persons: increased susceptibility to apoptosis of CD4 and CD8 T cells correlates with lymphocyte activation and with disease progression. J Immunol 156, 3509–3520 (1996).

48. N. Selliah, T. H. Finkel, Biochemical mechanisms of HIV induced T cell apoptosis. Cell Death Differ 8, 127–136 (2001).

49. J. A. Akulian et al., High-quality CMV-specific CD4+ memory is enriched in the lung allograft and is associated with mucosal viral control. Am J Transplant 13, 146–156 (2013).

50. C. Monaco, J. Nanchahal, P. Taylor, M. Feldmann, Anti-TNF therapy: past, present and future. Int Immunol 27, 55–62 (2015).

51. E. J. Giamarellos-Bourboulis et al., Complex Immune Dysregulation in COVID-19 Patients with Severe Respiratory Failure. Cell Host Microbe 27, 992–1000 e1003 (2020).

52. D. M. Del Valle et al., An inflammatory cytokine signature predicts COVID-19 severity and survival. Nat Med 26, 1636–1643 (2020).

53. P. C. Robinson et al., The Potential for Repurposing Anti-TNF as a Therapy for the Treatment of COVID-19. Med (N Y) 1, 90–102 (2020).

54. E. Valenzi et al., Single-cell analysis reveals fibroblast heterogeneity and myofibroblasts in systemic sclerosis-associated interstitial lung disease. Ann Rheum Dis 78, 1379–1387 (2019).

55. M. Stoeckius et al., Cell Hashing with barcoded antibodies enables multiplexing and doublet detection for single cell genomics. Genome Biol 19, 224 (2018).

56. T. Stuart et al., Comprehensive Integration of Single-Cell Data. Cell 177, 1888–1902 e1821 (2019).

57. A. Ferguson, K. Chen, Analysis of Transcriptional Profiling of Immune Cells at the Single-Cell Level. Methods Mol Biol 2111, 47–57 (2020).

58. A. Liberzon et al., Molecular signatures database (MSigDB) 3.0. Bioinformatics 27, 1739–1740 (2011).

